# Dispersal of PRC1 condensates disrupts polycomb chromatin domains and loops

**DOI:** 10.1101/2023.02.22.529502

**Authors:** Iain Williamson, Shelagh Boyle, Graeme Grimes, Elias Friman, Wendy A. Bickmore

## Abstract

Polycomb-repressive complex 1 (PRC1) has a strong influence on 3D genome organization, mediating local chromatin compaction as well as localized and chromosome-wide clustering of target loci. Several subunits of PRC1 have been shown to have the capacity to form biomolecular condensates through liquid-liquid phase separation *in vitro* and when tagged and over-expressed in cells. Here, we use 1,6-hexandiol, which disrupts liquid-like condensates, to examine the role of endogenous PRC1 biomolecular condensates on local and chromosome-wide clustering of PRC1-bound loci. Using imaging and chromatin immunoprecipitation combined with deep sequencing analyses, we show that PRC1-mediated chromatin compaction and clustering of targeted genomic loci – at megabase and tens of megabase length scales – can be reversibly disrupted by the addition and subsequent removal of 1,6-hexandiol to mouse embryonic stem cells. Decompaction and dispersal of polycomb domains and clusters cannot be solely attributable to the reduction of PRC1 binding following 1,6-hexandiol treatment as the addition of 2,5-hexandiol has similar effects on binding despite this alcohol not perturbing PRC1-mediated 3D clustering, at least at the sub-megabase and megabase scales. These results suggest that weak hydrophobic interactions between PRC1 molecules, characteristic of liquid-liquid phase separation, have a role in polycomb-mediated genome organization.

## Introduction

The spatial organisation of biochemical reactions in the cell nucleus is fundamental to genome regulation. The concentration of nucleic acids and proteins in membrane-less compartments, or condensates, can increase the kinetics of, and modulate the outcomes of, biochemical reactions (Holehouse and Pappu, 2018). Condensates have been implicated in the regulation of a multitude of nuclear functions including; DNA repair, replication, transcription, RNA processing and epigenetic regulation (Banani et al., 2017). Many biophysical mechanisms can result in the formation of condensates, but one receiving a lot of attention currently is liquid-liquid phase separation (LLPS). Proteins that tend to undergo LLPS often have regions that participate in multivalent weak interactions, that are intrinsically disordered, or that favour oligomerisation (Banani et al., 2017).

Chromatin can act as a scaffold for phase separation and chromatin-binding proteins have been implicated in the formation of phase-separated chromatin compartments (Bajpai et al., 2021). Interest in this area was propelled by observations of the LLPS properties of the chromobox (Cbx) containing heterochromatin protein 1α (HP1α)/Cbx5 (Larson et al., 2017; Strom et al., 2017). HP1 proteins recognise and bind to histone H3 tri-methylated at lysine 9 (H3K9me3) through their Cbx domain. However, the role of LLPS in the compaction of H3K9me3-marked constitutive heterochromatin has been questioned and instead it has been suggested that the ability of HP1α to bridge between H3K9me3-modified nucleosomes is sufficient to drive the compaction of constitutive heterochromatin (Erdel et al., 2020).

Cbx-domain containing proteins are also important in the formation of facultative heterochromatin. Cbx2,4,6,7 and 8 are components of the canonical polycomb-repressive complex 1 (cPRC1) and are responsible for the targeting of this complex to H3K27me3-modified chromatin, deposited by the PRC2 complex. Cbx2 has been shown to be able to phase separate in vitro (Plys et al., 2019) and when ectopically expressed in mammalian cells (Tatavosian et al., 2019), though its behaviour when induced to oligomerise in cells is more complex (Eeftens et al.,2021). In addition, other components of PRC1 have the ability to form condensates under those conditions and indeed the polyhomeotic components of cPRC1 are of particular interest because their sterile alpha motif (SAM)-domain is known to drive oligomerisation (Isono et al., 2013), can phase separate in vitro, form condensates in cells (Seif et al., 2020; Eeftens et al., 2021) and can compact chromatin (Kundu et al., 2017). Most of these studies have analysed the properties of cPRC1 components in vitro, or when tagged and over-expressed in cells. There has been little detailed analysis of the role of weak multivalent interactions in modulating the local and long-range chromatin conformation at target loci of the endogenous cPRC1 complex.

Both imaging and chromosome conformation analyses have shown that in mouse embryonic stem cells (mESCs) polycomb acts not only to locally compact target loci (Eskeland et al. 2010; Kundu et al. 2017; Boyle et al., 2020) but also to bring together distant target loci (Isono et al. 2013; Schoenfelder et al. 2015; Vieux-Rochas et al. 2015; Kundu et al. 2017; Boyle et al., 2020). This clustering brings multiple polycomb target loci close together in nuclear space and occurs over genomic distances of 10s to 100s of Mb – far beyond the scale of topologically associating domains (TADs) (Boyle et al., 2020). Consistent with this, long-range polycomb associations do not depend on cohesin (Rhodes et al., 2020).

Here we investigate the role of multivalent weak interactions in the ability of endogenous polycomb complexes to direct 3D chromatin organisation in mESCs. Using DNA fluorescence in situ hybridisation (DNA-FISH), we assay chromatin compaction of single large (> 100 kb) polycomb-bound domains and clustering of short-(100 kb – 2 Mb) and long-range (10s of megabases) polycomb target loci in cells treated with two aliphatic alcohols, one with and one without the capacity to disrupt biomolecular condensates, as well as in cells allowed to recover after alcohol treatment. We show that the perturbation of cPRC1 condensates reversibly disrupts chromatin compaction and clustering of polycomb target loci. Using chromatin immunoprecipitation followed by deep sequencing (ChIP-seq) we show that these effects cannot be solely explained by the loss of cPRC1-binding at target genomic loci. We suggest that multivalent weak interactions between polycomb complexes drives the ability of cPRC1 to modulate 3D genome organisation.

## Results

### Optimizing hexanediol treatment of mESCs

A significant factor hindering the investigation of phase separation in cells is the limited tools available to modulate the molecular interactions between endogenous macromolecules that drive these transitions. Many biomolecular condensates and nuclear bodies are sensitive to the aliphatic alcohol 1,6-hexanediol (1,6-HD), which disrupts weak multivalent hydrophobic interactions, and 1,6-HD has been used to discriminate liquid-like condensates from those with more solid or gel-like properties (Kroschwald et al. 2017). The aliphatic alcohol 2,5-hexanediol (2,5-HD) does not show this property (Lin et al., 2016; Kroschwald et al. 2017).

The addition of 1,6-HD to several human and mouse cell lines at concentrations ranging from 1.5% to over 10%, and with treatments lasting from less than a minute to thirty minutes, has been used to disrupt nuclear biomolecular condensates, including those implicated in chromatin states. However, some results have been conflicting and indicative of irreversible changes in chromatin and nuclear organisation (Itoh et al., 2021; Liu et al., 2021; Ulianov et al., 2021). We therefore tested a range of conditions for treating mESCs with 1,6-HD (2%, 5% and 10% concentrations for 5, 10 and 15-minute durations prior to fixation) to minimise harmful side-effects and prevent excessive cell death. For mESCs growing in tissue-culture flasks and on microscope glass slides we found that both 5% and 10% concentrations of 1,6-HD caused excessive cell-death. Visually, cells grown in flasks treated with 2% 1,6-HD or 2,5-HD appeared normal up to 10 minutes treatment duration with minimal cell-death, but for cells growing on glass slides optimal treatment duration was reduced to 5 minutes (Fig. 1A; Supplemental Fig. S1A). The morphology of nuclei in mESCs treated with 2% 1.6-HD or 2,5-HD for 5 minutes appeared to be no different to untreated cells and cells allowed to recover for over one hour post-treatment, with chromocentres readily apparent (Fig. 1B), and nuclear size was also unaffected by HD treatment (Fig. 1C; Supplemental Fig. S1B; Supplemental Table S1). Therefore, for all subsequent experiments we treated cells with 2% 1,6-HD or 2,5-HD for 5 minutes, which corresponds well with the optimal parameters for 1,6-HD treatment of mESCs reported in a recent study (Liu et al., 2021).

**Figure 1.**
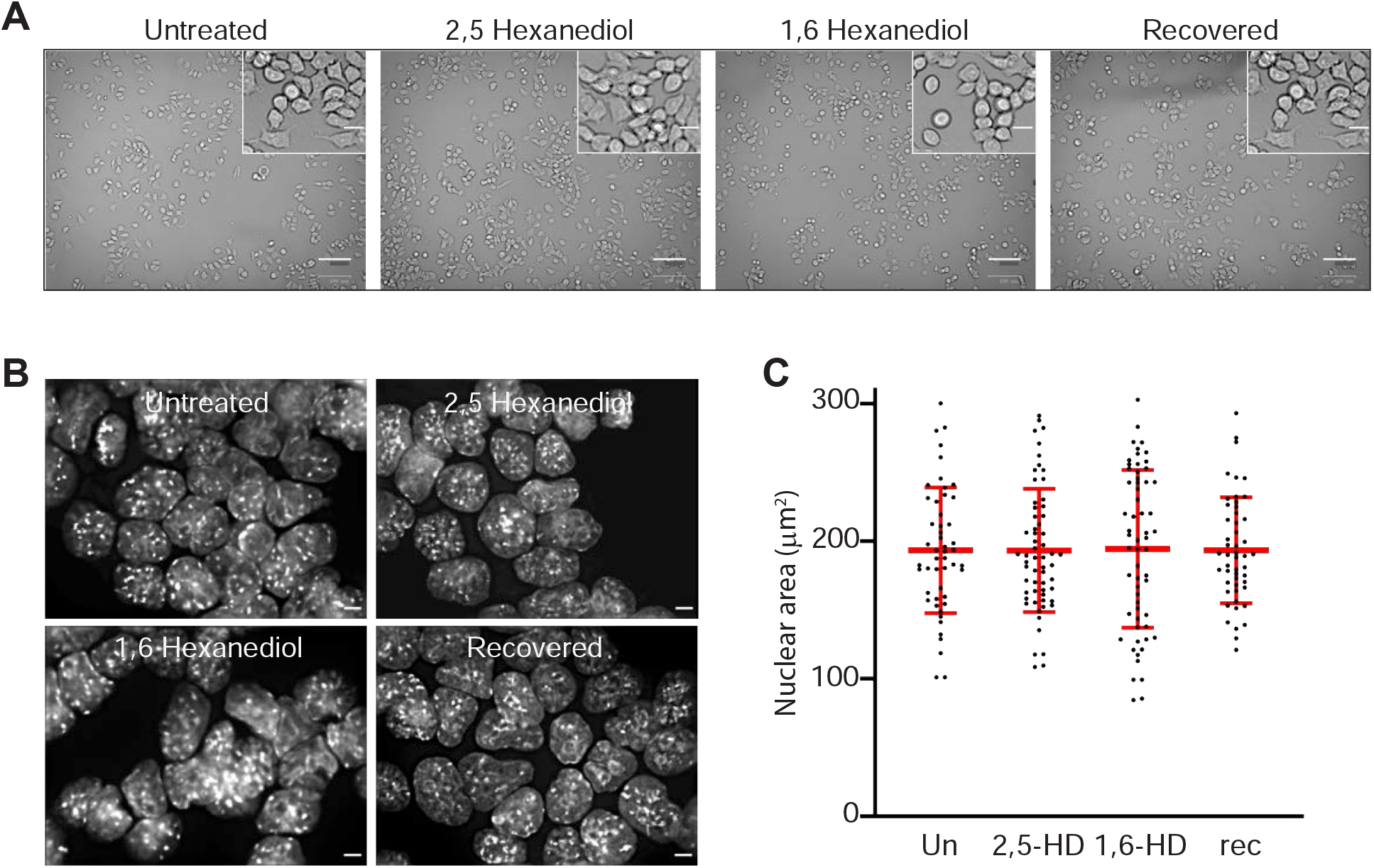
Optimal hexandiol application conditions for mESCs are 2% concentration for 5 minutes. (*A*) Representative phase contrast images of untreated, 2,5-HD-treated, 1,6-HD-treated, and recovered mESCs. Scale bars: 100 μm. Inset scale bars: 20 μm. (*B*) Representative images of DAPI-stained untreated, 2,5-HD-treated, 1,6-HD-treated, and recovered mESC nuclei. Scale bars: 5 μm. (*C*) Scatter plot showing the nuclear areas (μm^2^) of untreated, 2,5-HD-treated, 1,6-HD-treated, and recovered mESCs (black dots). Red bars show the means and standard deviations. Data sets were tested for differences using the Unpaired T-test with Welch’s correction. Results from a biological replicate experiment are shown in Supplemental Fig. S1 and the data for these figures are in Supplemental Table S1.

### 1,6-HD but not 2,5-HD treatment reversibly decompacts Hox clusters

Components of the polycomb repressive complex 1 (PRC1) have been shown to be capable of phase separating in vitro, or when ectopically expressed (Plys et al., 2019; Tatavosian et al., 2019; Seif et al., 2020). We therefore sought to determine whether the chromatin changes induced by cPRC1 binding at endogenous PRC1 target loci in mESCs are sensitive to disruption by 1,6-HD.

The four paralogous Hox loci, containing densely packed arrays of homeobox genes, are the largest polycomb targets in mESCs, forming compact chromatin domains of over 100kb in size that are dependent on PRC1 and PRC2 (Eskeland et al., 2010; Williamson et al., 2014; Kundu et al. 2017; Boyle et al., 2020). As an example, Hi-C analysis identifies highly enriched interactions across the HoxD region in mESCs, including the neighbouring *Evx2* gene, that correspond exactly with the extent of H3K27me3 and Ring1B (PRC1 subunit) occupancy (Fig. 2A, upper heatmap and ChIP-seq tracks). This enriched interaction domain is lost in mESCs without functioning PRC1 (*Ring1B*^*-/*-^) (Fig. 2A, lower heatmap).

**Figure 2.**
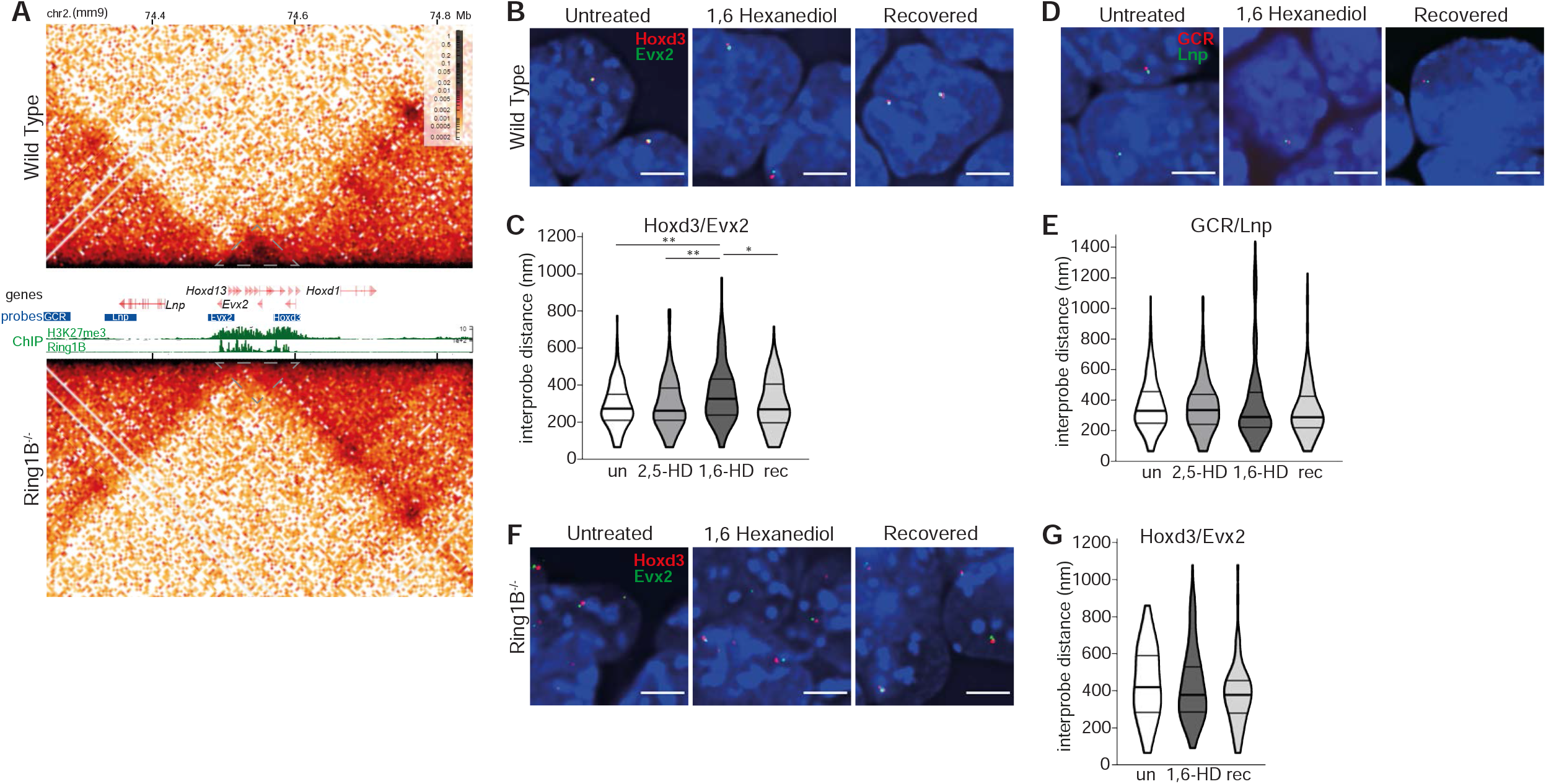
Reversible chromatin decompaction at the HoxD cluster following 1,6-HD treatment. (*A*) Hi-C maps (5kb resolution) of the HoxD cluster from wild-type (top: *Ring1B*^*+/+*^) and (lower) *Ring1B*^*− /−*^ mESCs. Genome co-ordinates on chromosome 2 (Mb) from the mm9 assembly of the mouse genome. Genes, fosmid probe binding locations, H3K27me3 and RING1B ChIP-seq profiles are shown between the two Hi-C maps. Grey dotted triangles indicate the extent of the HoxD locus (data are from Boyle et al., 2020). (*B* and *D*) Representative 3D-FISH images for polycomb target (*B*) and non-polycomb (*D*) target loci at HoxD in untreated, 1,6-HD-treated and recovered wild-type mESCs. Scale bars: 5 μm. (*C* and *E*) Violin plots show the interprobe distances (nm) for probe pairs used in *B* and *D* in untreated, 2,5-HD-treated, 1,6-HD-treated, and recovered wild-type cells. ∗P ≤ 0.05 and > 0.01; ∗∗P ≤ 0.01; Mann Whitney test. Data for a biological replicate are shown in Supplemental Fig. S2 and data are summarised in Supplemental Table S2. (*F* and *G*) As in *B* and *C* but using *Ring1B*^*− /−*^ mESCs.

We have previously shown by 3D FISH that loss of PRC2 or PRC1 results in a significant visible increase in spatial distance between the 3′ and 5′ ends of the HoxD cluster in nuclei, indicative of localised chromatin decompaction, whereas the non-polycomb target region upstream of HoxD (limb enhancers GCR to *Lnp*) is unaffected (Eskeland et al., 2010; Williamson et al., 2014). To determine whether polycomb-mediated chromatin compaction is dependent on weak multivalent interactions, we measured the inter-probe distances across HoxD (probes Evx2 and Hoxd3) and the control region (between GCR and Lnp) (Fig. 2A, B, D) in cells treated with 1,6-HD and 2,5-HD, compared with untreated cells and cells allowed to recover in normal media for at least one hour post 1,6-HD treatment. Inter-probe distances between the 3′ and 5′ ends of HoxD significantly increased upon treatment with 1,6-HD, with the distances in recovered cells returning to levels similar to those seen in untreated cells (Fig. 2B, C; Supplemental Fig. S2A; Supplemental Table S2). In contrast, there was no significant effect on interprobe distances at the nearby control non-polycomb target region (Fig. 2E; Supplemental Fig. S2B; Supplemental Table S2). Treatment with 2,5-HD had no significant effect at either locus. 1,6-HD also resulted in increased distances separating the ends of HoxB, with the cluster also returning to a more compact conformation in recovered cells (Supplemental Fig. S2C; Supplemental Table S2).

The influence of 1,6-HD on chromatin compaction at HoxB and HoxD, but not at a nearby control locus, strongly implicates a specific effect of 1,6-HD on polycomb marked chromatin. To investigate this further, we carried out the same analysis on *Ring1B*^*-/-*^ mESCs (Fig. 2F). As we have shown previously (Eskeland et al., 2010; Boyle et al., 2020), the HoxD cluster is in a decompact conformation in *Ring1B*^*-/-*^ mESCs compared to wild-type cells (Fig. 2G; Supplemental Table S2). Treatment of *Ring1B*^*-/-*^ mESCs with 1,6-HD had no further effect on interprobe distances measured across the HoxD cluster (Fig. 2G; Supplemental Table S2).

These results suggest that disruption of weak multivalent interactions causes a specific and reversible loss of chromatin compaction at Hox loci that is dependent on cPRC1.

### Reversible disruption of dispersed homeobox gene clusters by 1,6-HD

In addition to causing local chromatin compaction, PRC1 has also been implicated in bringing together more distal polycomb targets (Isono et al. 2013; Schoenfelder et al. 2015; Vieux-Rochas et al. 2015; Kundu et al. 2017; Boyle et al., 2020). Whereas the *Hox* genes are densely packed within the ∼100kb of each of the Hox loci, other clusters of vertebrate genes encoding homeodomain transcription factors are more dispersed. An example of the latter are the *Irx* gene clusters encoding the Iroquois transcription factors (Peters et al., 2000). *Irx3, Irx5* and *Irx6* are spread across 900kb of mouse chromosome 8, albeit within the same large (∼2Mb) TAD (Sobreira et al., 2021). In mESCs all three genes are polycomb targets and form polycomb (RING1B)-dependent associations with each other in Hi-C (Fig. 3A) (Boyle *et al*., 2020) and micro-C data, and these associations are unaffected by the loss of either CTCF or cohesin (Supplemental Fig. S3A) (Hsieh et al., 2020; Hsieh et al., 2022). This association is reflected in the tight nuclear co-localisation of *Irx3, Irx5* and *Irx6* as determined by DNA-FISH (Fig. 3B). Two out of three of *Irx3, Irx5* and *Irx6* are defined as clustered (<200nm separation) at a quarter of measured alleles (Fig. 3C; Supplemental Fig. S3B, C; Supplemental Table S3). Indeed, for a large proportion of alleles all three loci cluster within 400 nm of each other (Fig. 3D). All three loci were separated by >400nm in <20% of alleles.

**Figure 3.**
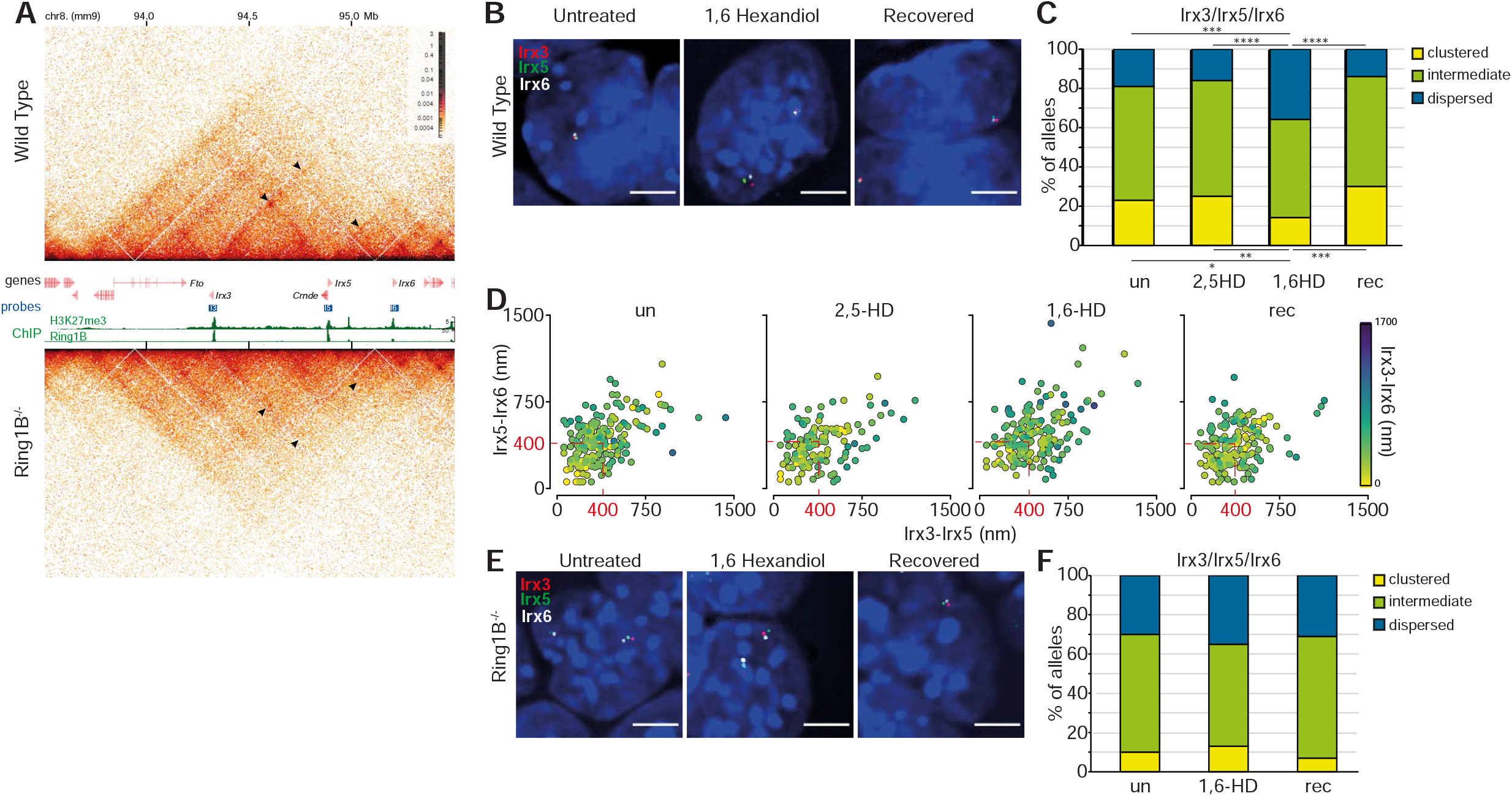
Reversible disruption of *Irx* gene clustering by 1,6-HD. (*A*) Hi-C maps of the TAD on chromosome 8 (mm9 co-ordinates in Mb) containing *Irx3, 5* and *6* from (upper) wild-type (*Ring1B*^*+/+*^) and (lower) *Ring1B*^*− /−*^ mESCs at 5kb resolution. Genes, fosmid probe binding locations, H3K27me3 and RING1B ChIP-seq profiles are shown between the two Hi-C maps (data are from Boyle *et al*., 2020). Black arrowheads indicate enriched interactions between *Irx3, 5* and *6* in wild-type mESCs and the reduced interactions in *Ring1B*^*− /−*^ cells. (*B*) Representative 3D-FISH images of *Irx3, 5* and *6* loci in untreated, 1,6-HD-treated and recovered wild type mESCs. Scale bar: 5 μm (*C*) Bar plots providing categorical analysis of the spatial relationship of *Irx3, 5* and 6 in untreated, 2,5-HD-treated, 1,6-HD-treated and recovered wild-type cells. Categories are: clustered, at least two of the three loci < 200 nm apart; intermediate, at least two of the three loci between 200 – 399 nm apart; dispersed, all three loci ≥ 400 nm apart. Differences in clustering and dispersal identified using Fisher’s Exact test; ∗P ≤ 0.05 and > 0.01, ∗∗P ≤ 0.01, ∗∗∗P ≤ 0.001, ∗∗∗∗P ≤ 0.0001. (*D*) Scatter plots showing interprobe distances between each of the two probe pairs indicated on *x* and *y* axes with the separation between the third pair indicated by the color (in the color bar) in untreated, 2,5-HD-treated, 1,6-HD-treated and recovered wild-type mESCs. Dashed red box indicates non-dispersed alleles with at least two sets of interprobe distances < 400 nm. (*E* and *F*) As for *B* and *C*, but for *Ring1B*^*-/-*^ cells. Scatter plots for those data are in Supplemental Fig. S3, as are data for a biological replicate. All data are summarised in Supplemental Table S3.

These proportions were not significantly different in 2,5-HD treated cells. However, *Irx* clustering was significantly disrupted in 1,6-HD-treated mESCs cells (Fig. 3C, D; Supplemental Fig. S3B, C; Supplemental Table S3) and was restored after 1,6-HD removal (recovered). *Irx* clustering was low (7-13%) in *Ring1B*^*-/-*^ mESCs and most alleles contained *Irx3, 5* and *6* in more dispersed configurations. There was no further dispersal of these three loci in 1,6-HD treated *Ring1B*^*-/-*^ cells (Fig. 3E, F; Supplemental Fig. S3D; Supplemental Table S3). Therefore, the effect of 1,6-HD can be attributed to polycomb function.

### Reversible disruption of long-range inter-TAD polycomb interactions by 1,6-HD

Interactions between distal polycomb targets can occur over large genomic distances that span beyond a single TAD (Boyle et al., 2020). As an example, *Shh* located within a 900kb TAD, is one of only three polycomb target genes across a 2Mb region of chromosome 5 in mESCs, the other two genes – *En2* and *Mnx1* – being in the flanking TADs (Fig. 4A). Hi-C (Fig. 4A) and Micro-C (Supplemental Fig. S4A) (Hsieh et al., 2020; Hsieh et al., 2022) identifiy enriched interactions between *En2, Shh* and *Mnx1* that are lost in Ring1B null cells (Fig. 4A) (Boyle et al., 2020) but that are unaffected by the loss of CTCF or RAD21 (Supplemental Fig. S4A). Spatial proximity between all three polycomb target genes could also be detected, which was significantly reduced in Ring1B null cells, or when one of the three genes are expressed *in vivo* (Boyle et al., 2020). We performed 3D-FISH, using probes covering polycomb target and adjacent non-target loci (Boyle et al., 2020) (Fig. 4A), on untreated, 1,6-HD- and 2,5-HD-treated, and recovered mESCs (Fig. 4B). As was the case for the intra-TAD polycomb associations at the *Irx* locus, clustering of *Shh, En2*, and *Mnx1* was significantly reduced by 1,6-HD treatment, and restored after 1,6-HD removal (rec); whereas 2,5-HD treatment had no effect (Fig. 4C, D; Supplemental Fig. S4B, C; Supplemental Table S4). Clustered and dispersed ratios remained unchanged at the adjacent non-polycomb target loci (*Cnpy1*, SFPE1, *Ube3c*) across the various treatments, with high proportions of dispersed alleles (Fig. 4E Supplemental Fig. S4D, E; Supplemental Table S5). Addition of 1,6-HD had no significant effect on the spatial positioning of the three polycomb target loci in *Ring1B*^*-/-*^ cells, with no further dispersal of *En2, Shh* and *Mnx1* (Fig. 4F; Supplemental Fig. S4F and Table S4).

**Figure 4.**
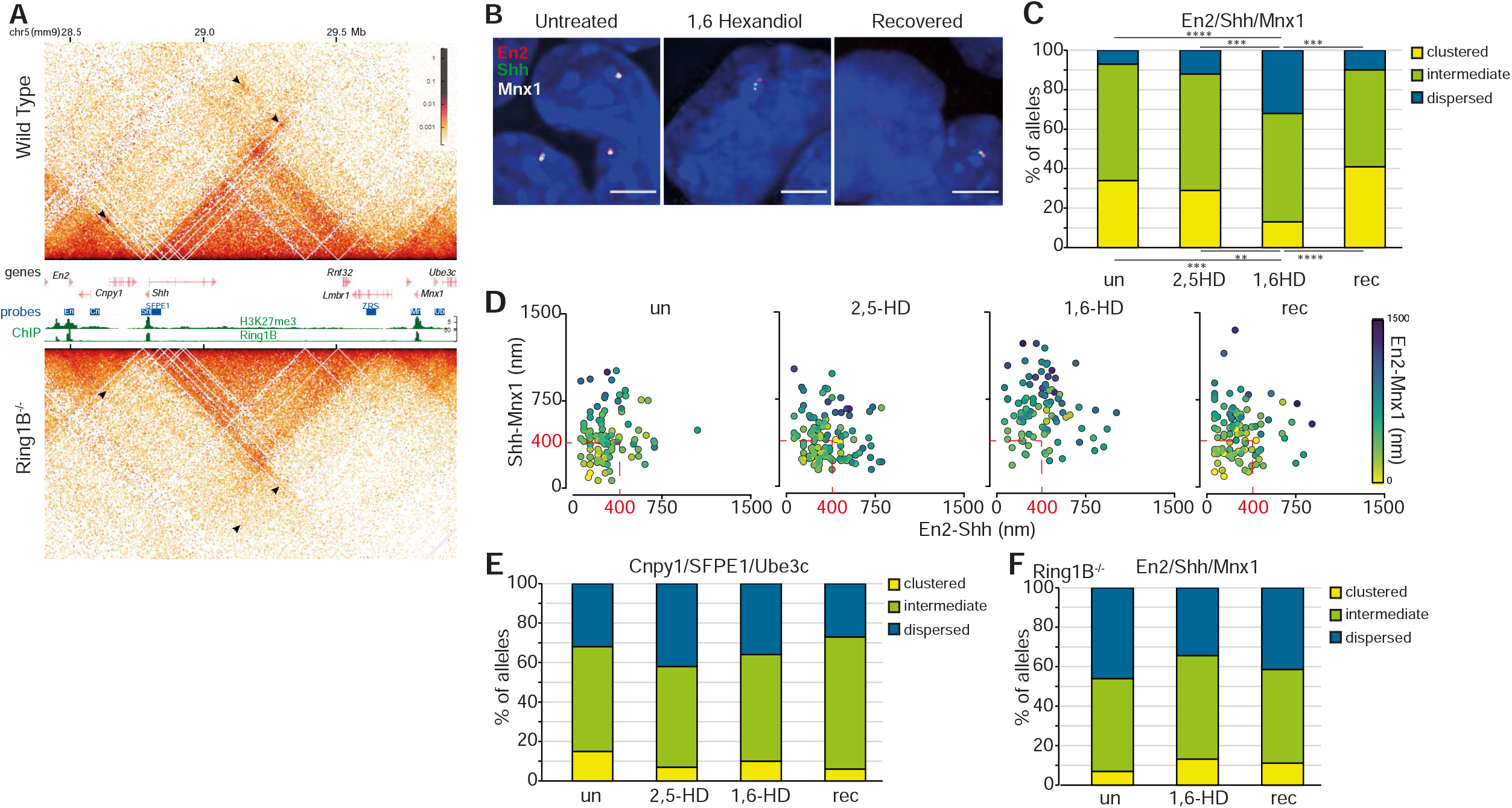
Reversible disruption of inter-TAD polycomb clustering by 1,6-HD. (*A*) Hi-C maps of the three adjacent TADs on chromosome 5 (mm9) containing *En2, Shh*, and *Mnx1* polycomb targets from Wild-Type (*Ring1B*^*+/+*^) and *Ring1B*^*− /−*^ mESCs at 5kb resolution. Genes, fosmid probe binding locations, H3K27me3, and RING1B ChIP-seq profiles are shown between the two Hi-C maps (data from Boyle et al., 2020). Black arrowheads indicate enriched interactions between the three polycomb target genes in Wild-Type cells and reduced interactions in *Ring1B*^*− /−*^ cells. (*B*) Representative images of 3D-FISH for *En2, Shh*, and *Mnx1* in untreated, 1,6-HD-treated, and recovered wild type cells. (*C* and *E*) Bar plots providing categorical analysis of the spatial location of each of the three polycomb (*C*) and non-polycomb (*E*) probes shown in *A* relative to each other in untreated, 2,5-HD-treated, 1,6-HD-treated and recovered wild type cells. Categories are as in Figure 3. Differences in clustering and dispersal identified using Fisher’s Exact test; ∗∗P ≤ 0.01, ∗∗∗P ≤ 0.001, ∗∗∗∗P ≤ 0.0001. (*D*) Scatter plots depicting the interprobe distances between each of the two fosmid probe pairs indicated on *x* and *y* axes with the separation between the third pair indicated by the color in the color bar for polycomb target loci in Wild-Type cells. Dashed red box indicates non-dispersed alleles with at least two sets of interprobe distances < 400 nm. (*F*) As in C but for untreated, 1,6-HD-treated and recovered *Ring1B*^*-/-*^ cells. Data for a biological replicate are in Supplemental Fig. S4 and data are summarised in Supplemental Tables S4 and S5.

There is a strong interaction between the TAD boundaries involving CTCF sites adjacent to *Shh* and ZRS limb enhancer (Williamson et al., 2016, 2019). To determine if the loss of spatial proximity between *Shh, En2*, and *Mnx1* was influenced by a general perturbation of local chromatin organisation affecting non-polycomb associations we compared spatial distances between probes covering *Shh* and ZRS and their adjacent CTCF sites. No significant differences in spatial proximity between these two loci were identified in 2,5-HD, 1,6-HD and recovered cells compared with untreated cells (Supplemental Fig. S4G). Therefore, disruption of PRC1 condensates causes the specific dispersal of polycomb dependent associations between three polycomb target genes located across three adjacent TADs over a 2Mb distance.

### Hexandiol treatment reversibly perturbs chromosome-wide clustering of polycomb targets

Previously, we showed that cPRC1 has a constraining influence on chromosomal architecture through extreme long-range, chromosome-wide, clustering of polycomb target loci (Boyle et al., 2020). To determine whether cPRC1 associations at this length scale are also sensitive to disruption by 1,6-HD, we used two sets of oligo probe pools, one labelling polycomb target loci (PcG+), the other intervening non-polycomb target regions (PcG-) covering 28 discrete loci along 51Mb of chromosome 2 (Fig. 5A).

**Figure 5.**
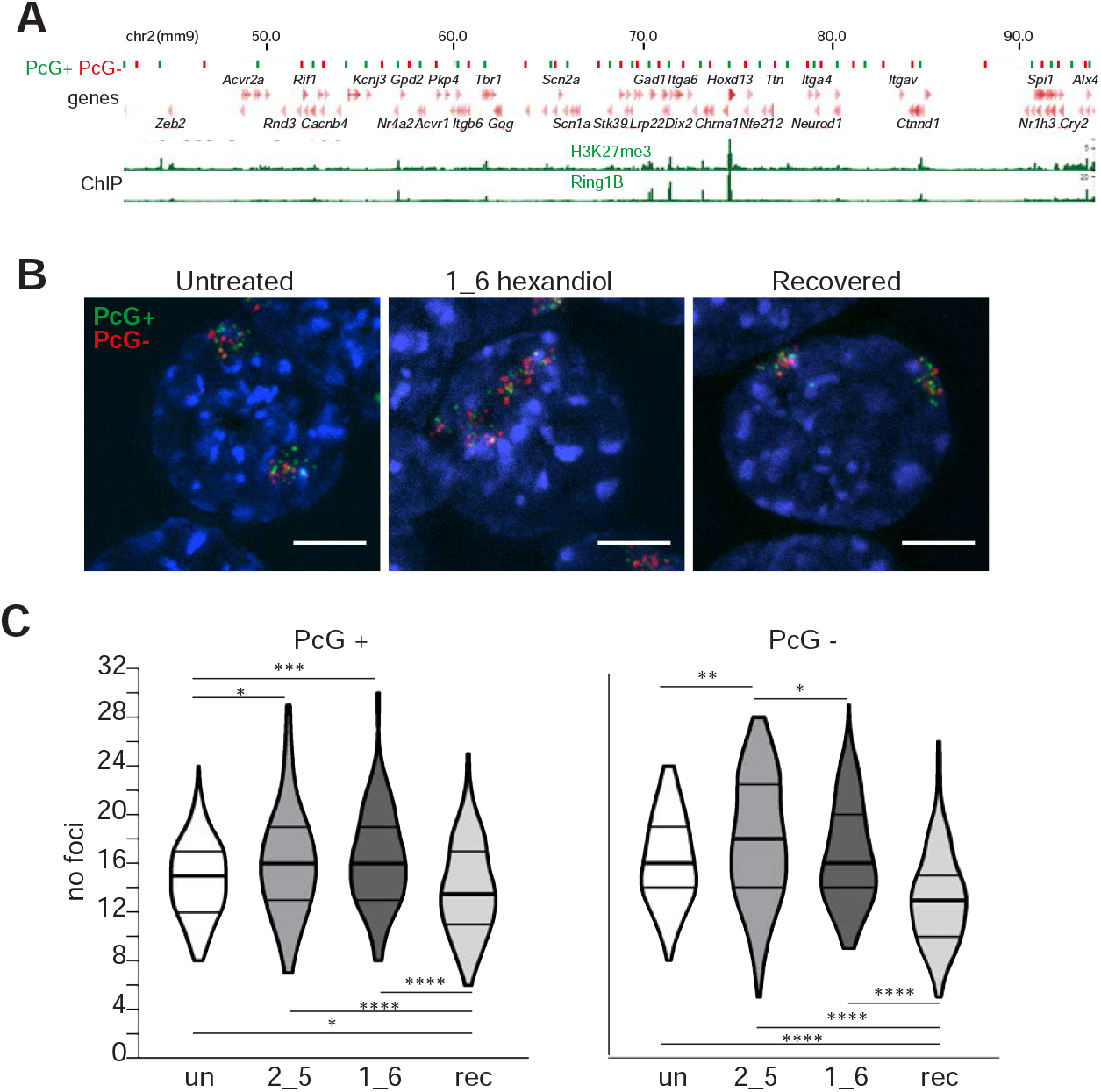
Reversible loss of PRC1 dependent clustering of polycomb targets across chromosome 2 following hexandiol treatment. (*A*) Ideogram of chromosome 2 indicating the location of the oligonucleotide probes used in *B* and *C* and zoomed in browser tracks of RING1B and H3K27me3 ChIP-seq from wild-type mESCs (Boyle *et al*., 2020). Polycomb-positive (PcG+) and polycomb-negative (PcG−) probe locations are represented as green and red bars, respectively. Genomic co-ordinates (Mb) are for the mm9 genome assembly. (*B*) Representative 3D-FISH images of untreated, 1,6-HD-treated, and recovered cells hybridized with the chromosome 2 PcG+ (green; 6FAM) and PcG− (red; ATT0594) probe pools. Scale bar: 5 μm. (*C*) Violin plots depicting the number of discrete foci in untreated, 2,5-HD-treated, 1,6-HD-treated and recovered cells for PcG+ and PcG− probe pools. ∗P≤0.05 and > 0.01; ∗∗P≤0.01; ∗∗∗P ≤ 0.001, ∗∗∗∗P ≤ 0.0001; Mann Whitney test. Data are summarised in Supplemental Table S6.

As in the absence of PRC1, the addition of 1,6-HD caused significantly reduced clustering of polycomb target loci, and not for the intervening non-polycomb regions (Fig. 5B, C; Supplemental Table S6). 2,5-HD treatment caused a more subtle disruption of polycomb target clustering but the addition of this alcohol also reduced clustering of the non-polycomb intervening loci, suggesting a chromosome-wide effect of this alcohol (Fig. 5C; Supplemental Table S6). Surprisingly, enhanced clustering of both PcG+ and PcG-probe pools was detected in recovered cells, compared with untreated cells, (Fig. 5B, C; Table S6). This suggests a persistent effect of 1,6-HD at the chromosome territory level.

### Perturbation of chromatin organisation by 1,6-HD is not simply the consequence of loss of PRC1 binding

The influence of cPRC1 on multiple levels of chromatin organisation, from individual chromatin domains such as the Hox clusters, to clustering of polycomb target loci over various length scales, has been demonstrated by the perturbation of these structures in *Ring1B*^*-/-*^ cells (Boyle et al., 2020). To determine if the specific effects of 1,6-HD, and not 2,5-HD, on these same levels of 3D chromatin organisation could be attributed to the loss of PRC1 binding to chromatin following treatment, we performed calibrated ChIP-seq (cChIP-seq (Hu et al., 2015)) for Ring1B in mESCs treated with 1,6-HD, or 2,5-HD and in recovered cells.

Substantially reduced RING1B binding was detected genome-wide at target loci in cells treated with both 2,5-HD and 1,6-HD (Fig. 6A; Supplemental Fig. S5A). Ring1B binding was restored in recovered cells. This was also the case when examining specific loci (Fig. 6B, C; Supplemental Fig. S5B, C). We saw no evidence for preferential retention of Ring1B at specific sites in 1,6-HD- or 2,5-HD-treated cells.

**Figure 6.**
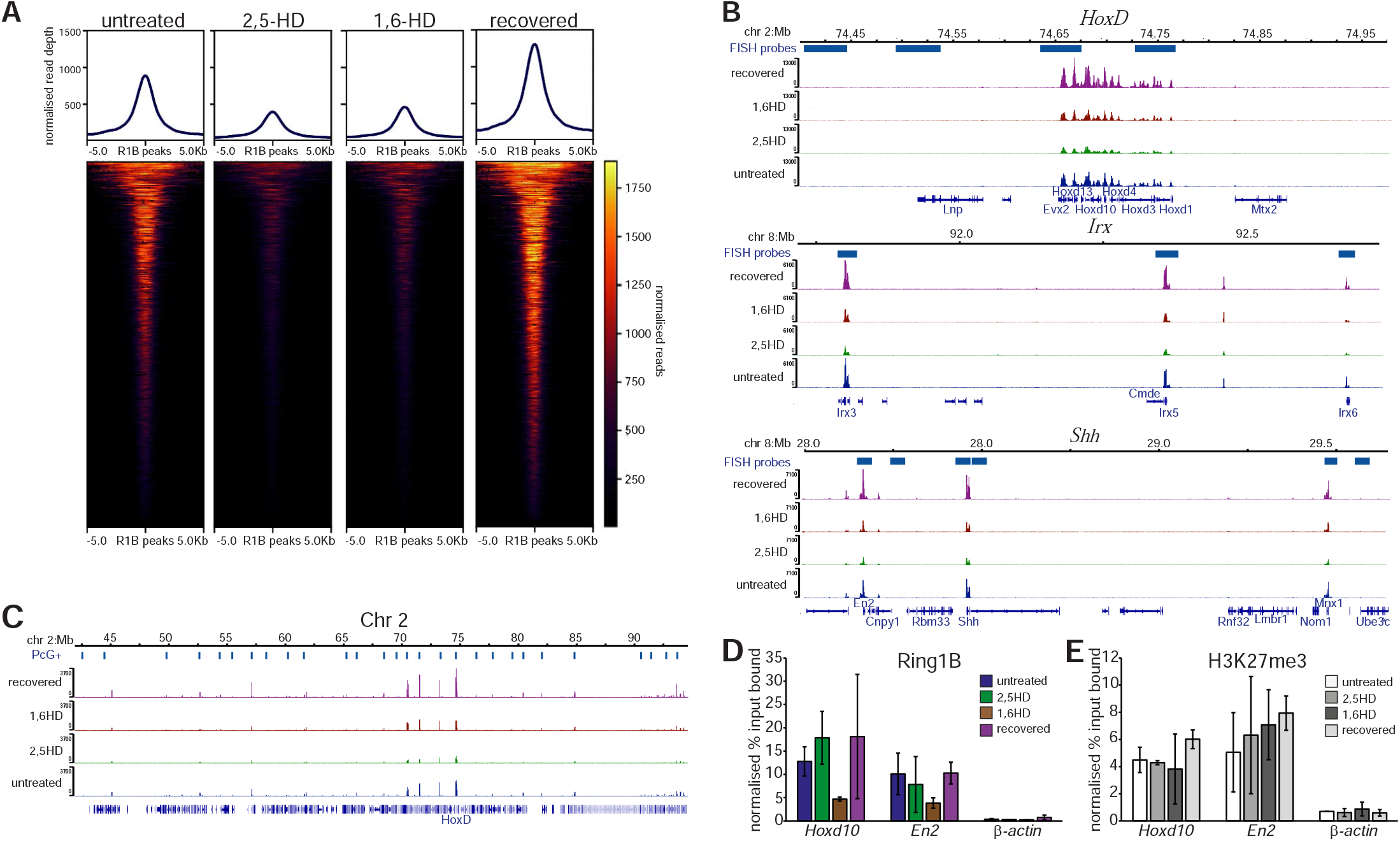
Reversible reduction of PRC1 binding to polycomb target loci following hexandiol treatment. (*A*) Heatmap representation (bottom panel) and average metaplots (top panel) of RING1B calibrated ChIP-seq signal distribution at merged RING1B peaks in untreated, 2,5-HD-treated, 1,6-HD-treated, and recovered wild-type mESCs. (*B*) UCSC Genome Browser track views of RING1B cChIP-seq data from untreated, 2,5-HD-treated, 1,6-HD-treated, and recovered mESCs at the HoxD (*top*), *Irx3, 5, 6* (*middle*), *En2, Shh*, and *Mnx1* (*bottom*) genomic regions. (*C*) As for *B* but for the 50 Mb region of chromosome 2 tiled with oligonucleotide probes. Biological replicate for these data are in Supplemental Fig. S5. (*D* and *E*) Normalised qPCR analysis of ChIP for RING1B (*D*) and H3K27me3 (*E*) at *Hoxd10, En2* and *β-actin* in untreated, 2,5-HD-treated, 1,6-HD-treated and recovered mESCs. Enrichment is shown as mean percent input bound ± s.e.m. over two biological replicates.

Quantitative ChIP-PCR confirmed the reduction of Ring1B occupancy after 1,6-HD treatment, and then its recovery, at *Hoxd10* and *En2* (Fig. 6D). This was not accompanied by any significant differences in levels of the PRC2-catalyzed histone mark H3K27me3 at these same loci (Fig. 6E). We conclude that the specific effects of 1,6-HD on chromatin compaction and long-range association between loci that are targets of cPRC1 – effects not seen with 2,5-HD – cannot simply be attributed to reduced RING1B chromatin binding, since that is affected by both alcohols.

## Discussion

Since PRC1 can mediate multivalent chromatin interactions that strongly influence local and long-range 3D genome organisation (Isono et al. 2013; Schoenfelder et al. 2015; Vieux-Rochas et al. 2015; Kundu et al. 2017; Boyle et al., 2020), and subunits of canonical PRC1 can undergo liquid-liquid phase separation (LLPS) (Plys et al., 2019; Tatavosian et al., 2019; Seif et al., 2020; Eeften et al., 2021), it is plausible that the 3D nuclear organisation of polycomb target loci in mammalian cells involves the formation of biomolecular condensates through processes such as phase separation. In support of this hypothesis, here we have shown by DNA-FISH in mESCs that PRC1-mediated compact chromatin domains, and the clustering of polycomb targets at length scales of many Mb, are reversibly perturbed by 1,6 hexandiol, but not by 2,5-HD. 1,6 hexandiol treatment had no effect on these loci in *Ring1B*^*-/-*^ mESCs showing that the sensitivity of these layers of 3D chromatin organisation to 1,6-HD is specific to PRC1 and is not due to a non-specific effect of this alcohol on chromatin organisation.

Using calibrated ChIP we show that 1,6-HD significantly reduces, but does not completely abolish, occupancy of the PRC1 subunit Ring1B at polycomb target loci genome-wide. However, reduced PRC1 occupancy also occurs in cells treated with 2,5-HD, a similar alcohol that does not have the same disruptive effects on chromatin compaction and polycomb target clustering as 1,6-HD. This implicates the loss of weak hydrophobic interactions between PRC1 molecules, rather than their reduced binding at target loci, as primarily responsible for the perturbations of polycomb domain compaction and distal interactions in the nucleus we have identified. Unlike conclusions drawn from ectopically expressed proteins (Eeften et al., 2021), our results suggest that chromatin compaction by PRC1 is dependent on condensate formation. The continued presence of H3K27me3 at polycomb loci after 1,6-HD treatment also excludes the possibility that a more generalised disruption of histone occupancy is responsible for the changes in 3D chromatin conformation.

We did find that extremely long-range (chromosome-wide) interactions between polycomb-sites were impacted by 2,5-HD treatment, though to a lesser extent than for 1,6HD (Fig. 5). Since 2,5-HD also seemed to decrease binding of PRC1 genome-wide, this result suggests that interactions of polycomb sites over very large length-scales, that are probably initiated through stochastic interactions in nuclear space, may rely, at least in part, on there being sufficiently high levels of PRC1 bound at target sites.

To preserve mESC morphology and cell viability, here we have only been able to induce very transient (5 minutes) disruption of polycomb-dependent 3D genome folding using 1,6-HD. Therefore, we are not able to determine the contribution that polycomb-mediated condensates may play in epigenetic mechanisms. Investigation of this interesting question awaits the development of less blunt tools for perturbing condensates in vivo.

## Materials and Methods

### Cell culture and alcohol treatments

Feeder-free mouse embryonic stem cells (mESCs) E14tg2A (129/Ola; *Ring1B*^*+/+*^) and the derivative line (*Ring1B*^*− /−*^) (Illingworth et al. 2015) were cultured at 37°C on 0.1% gelatin-coated (Sigma G1890) Corning flasks in GMEM BHK-21 (Gibco 21710-025) supplemented with 10% fetal calf serum (FCS; Sigma F-7524), 1000 U/mL LIF, nonessential amino acids (Gibco 11140-035), sodium pyruvate (Gibco 11360-039), 50 μM2-β-mercaptoethanol (Gibco 31350-010), and L-glutamine. For passaging, 60%–90% confluent flasks were washed with PBS, incubated for 2–3 minutes (mins) at room temperature in 0.05% (v/v) trypsin (Gibco 25300-054), and tapped to release. Trypsin was inactivated by adding 9 vol of ESC medium, and this mixture was repeatedly pipetted to obtain a single-cell suspension. ESCs were centrifuged, resuspended in ESC medium, and replated onto gelatin-coated flasks at a density of ∼4×10^4^ cells/cm^2^. For optimization of hexandiol experiments, ∼4×10^4^ cells/cm^2^ or ∼1×10^6^ cells were plated in standard medium, into flasks (for ∼48 hours) or on gelatin-coated SuperFrost plus microscopic glass slides (∼24 hours) respectively, before replacement with standard medium supplemented with 1,6 hexandiol at 2%, 5% and 10% concentrations (w/v) for 5, 10 or 15 mins. For subsequent experiments cells growing in flasks and on glass slide-plated cells were incubated in 2% 1,6 hexandiol (w/v) and 2,5 hexandiol (v/v) for 5 mins before harvesting or fixation. Medium containing 1,6 hexandiol was replaced with standard medium and cells were cultured for at least 1 hour (hr) before the harvesting or fixing of recovered cells. All centrifugation steps with live cells were performed at 330g for 4 min at room temperature. All ESC lines used in this study were routinely tested for mycoplasma.

### 3D DNA fluorescence in situ hybridization (DNA-FISH)

*Fixation:* Mouse ESCs grown on slides were fixed in 4% paraformaldehyde (pFA) for 10 minutes, permeabilized in PBS/0.5% Triton X100, dried, and then stored at − 80°C prior to hybridization. Slides were incubated in 100 μg/mL RNase A in 2× SSC for 1 hr at 37°C, washed briefly in 2× SSC, passed through an alcohol series, and air-dried. Slides were incubated for 5 min at 70°C, denatured in 70% formamide/2× SSC (pH 7.5) for 40 min at 80°C, cooled in 70% ethanol for 2 min on ice, and dehydrated by immersion in 90% ethanol for 2 min and 100% ethanol for 2 min prior to air drying.

### Hybridization and detection

For each slide hybridized with oligo probes, eight hundred nanograms of each fluorescently labeled oligonucleotide probe pool (2 μL; MyTags, Supplemental Tables S8, S9) (Boyle et al., 2020) were added to 26 μL of hybridization mix (50% formamide, 2× SSC, 1% Tween20, 10% dextran sulfate), denatured for 5 min at 70°C, and then snap-chilled on ice. For each slide hybridized with fosmid probes, 160-240 ng of biotin-, digoxigenin- and red-dUTP labelled (Alexa fluor 594-dUTP (Invitrogen)) fosmid probes (Supplemental Table S7), with 16-24 μg of mouse Cot1 DNA, and 10 μg of sonicated salmon sperm DNA were dried in a spin-vac and then reconstituted in 30 μL of hybridization mix. Probes were then denatured for 5 min at 80°C, reannealed for 15 min at 37°C.

Fosmid and oligonucleotide probes were hybridized to slides under a sealed coverslip overnight at 37°C. Slides were washed the next day four times for 3 min in 2× SSC at 45°C and four times for 3 min in 0.1× SSC at 60°C. Slides hybridized with directly labelled oligonucleotide probes were immediately stained with 4,6-diaminidino-2-phenylidole (DAPI) at 50 ng/mL, mounted in VectaShield (Vector Laboratories), and sealed with nail varnish. Slides hybridized with biotin- and dig-labelled fosmid probes were blocked in blocking buffer (4 x SSC, 5% Marvel) for 5 min. The following antibody dilutions were made: fluorescein anti-dig FAB fragments (Roche cat. no. 11207741910) 1:20, fluorescein anti-sheep 1:100 (Vector, cat. no. FI-6000)/ streptavadin Cy5 1:10 (Amersham, cat. no. PA45001, lot 17037668), biotinylated anti-avidin (Vector, cat. no. BA-0300, lot ZF-0415) 1:100, and streptavidin Cy5 1:10. Slides were incubated with antibody in a humidified chamber at 37°C for 30-60 min in the following order with 4X SSC/0.1% Tween 20 washes in between: fluorescein anti-dig, fluorescein anti-sheep/streptavidin Cy5, biotinylated anti-avidin, streptavidin Cy5 before staining and mounting with DAPI and vectashield respectively.

### Imaging and Image Analysis

Epifluorescent images were acquired using a Photometrics Coolsnap HQ2 CCD camera and a Zeiss AxioImager A1 fluorescence microscope with a plan apochromat 100× 1.4NA objective, a Lumen 200-W metal halide light source (Prior Scientific Instruments) and Chroma 89014ET single-excitation and emission filters (three-color FISH) or Chroma 89000ET single-excitation and emission filters (four-color FISH) (Chroma Technology Corp.) with the excitation and emission filters installed in Prior motorized filter wheels. A piezoelectrically driven objective mount (PIFOC model P-721, Physik Instrumente GmbH & Co.) was used to control movement in the z dimension. Hardware control, image capture, and analysis were performed using Volocity (Perkinelmer, Inc.) or Nis elements (Nikon). Images were deconvolved using a calculated point spread function with the constrained iterative algorithm of Volocity. Image analysis was carried out using the quantitation module.

### Hi-C and micro-C data

Hi-C data from *Ring1B*^*+/+*^ and *Ring1B*^*-/-*^ mouse embryonic stem cells (mESCs) were taken from Boyle et al, 2020. Micro-C WT mESC data were taken from Hsieh et al, 2020 and micro-C data from CTCF-AID and RAD21-AID IAA-treated ES cells were taken from Hsieh et al, 2022. Both Hi-C and micro-C data sets were visualised at 5 kb resolution using HiGlass (Kerpedjiev et al., 2018).

### Calibrated ChIP sequencing (ChIP-seq)

Calibrated ChIP-seq was carried out as previously described (Boyle *et al*., 2020). Briefly, trypsinized untreated, 2,5-HD, 1,6-HD and recovered mESCs (∼2 ×10^7^) were washed twice in PBS. After 10 minutes of formaldehyde (Calbiochem, cat. # 344198, final concentration 1%) fixation, stopped by 5 mins incubation with 125 mM of glycine at room temperature, cells were washed in PBS, pelleted, and stored at -80°C. Pellets were then thawed and combined with 1×10^6^ formaldehyde-fixed human MCF-7 cells (for downstream calibration of ChIP-seq data). Following lysis, cells were sonicated using a cooled Bioruptor (50 cycles, 1 minute cycles of 30s on/30s off on ‘high’ setting at 4°C). The sonicated extract was pre-cleared by centrifugation at 16,000g for 10 min at 4°C. The supernatant was transferred to a fresh tube and supplemented with BSA to a final concentration of 25 mg/mL. A sample of the chromatin was retained as an input reference.

Antibodies were precoupled to a 1/1 mixture of Protein A (Life Technologies 10001D) and Protein G (Life Technologies 1003D) Dynabeads at a ratio of 1 mg antibody per 30 mL of Dynabead suspension by rotation for 2 hr at 4°C. Cell equivalents (∼6.5×10^6^) of lysate were added to 7.5μg of anti-RING1B (Cell Signaling D22F2), 5 μg of anti-H3K27me3 (Cell Signaling C36B11), or 5 μg Rabbit IgG (Cell Signaling) respectively, and incubated overnight on a rotating wheel at 4°C. Following incubation, bead-associated immune complexes were washed sequentially with ChIP dilution buffer, wash buffer A, and wash buffer B, each for 10 min at 4°C on a rotating wheel, followed by two washes in TE buffer at room temperature. Chromatin was released by incubating the beads in 100 µL of elution buffer (0.1M NaHCO3, 1% SDS) for 15 min at 37°C, followed by the addition of 20 µg of RNase A and 6 µL of 2 M Tris (pH 6.8) and incubation for 1 h at 37°C and finally by the addition of 20 µg proteinase K and incubation overnight at 65°C to degrade proteins and reverse the cross-links. Dynabeads were removed using a magnetic rack and the chromatin purified using PCR purification columns (Qiagen) according to the manufacturer’s instructions.

Libraries (Ring1B ChIP and corresponding input samples only) were constructed using the NEBNext Ultra II DNA library preparation kit for Illumina according to the manufacturer’s instructions (NEB E7645S). To determine the number of PCR cycles required for amplification, one aliquot of library preparation from each sample were supplemented with EvaGreen so that amplification could be monitored by quantitative PCR on a BioRad C1000 Touch Thermal Cycler. To allow for sample multiplexing, PCRs were performed using index primers (NEBNext multiplex oligos for Illumina, set 1, E7335 and amplified to linear phase). Size selection purifications following the ligation and amplification PCR steps were performed with 0.9× reaction volumes of Agencourt AMPure XP beads (Beckman Coulter A63880). Purified libraries were combined as an 8-sample equimolar pool containing a combination of indexes from 1–12 and sequenced on an Illumina NextSeq on an Illumina NextSeq 2000 on a P2 flow cell (paired-end 75-bp reads).

### ChIP-seq analysis

Calibrated ChIP-seq data analysis was performed using the Nextflow (version 22.04.0) pipeline chip_quant_analysis.nf (https://gitbio.ens-lyon.fr/LBMC/Bernard/quantitative-nucleosome-analysis, 295ce7f5838d8fe08ad465232bdd178773a5f2c3) with the reference genome Mus musculus (GRCm38) and for internal calibration the Homo sapiens (GRCh38) genome. Briefly, Fastq files were processed with fastp (0.20.1, options --qualified_quality_phred 20 -- disable_length_filtering--detect_adapter_for_pe). The processed fastq files were mapped to a concatenated reference and calibration genome with bowtie2 (2.3.4.1, option --very-sensitive). Bam files were split into reads mapping exclusively to either the reference genome, or the calibration genome, using samtools (1.11), and awk. Mapped reads were deduplicated using picard (2.18.11, options MarkDuplicates VALIDATION_STRINGENCY=LENIENT REMOVE_DUPLICATES=true). The deduplicated bam files were converted to bigwig and bedgraph formats using deeptools (3.5.1). Normalization and generation of ratio coverage files were generated using software and scripts contained in the docker containers lbmc/chip_quant_r:<u>0.0.6</u> and biocontainers/danpos:v2.2.2_cv3 as described in in the paper https://www.biorxiv.org/content/10.1101/2022.03.18.484892v1.

Heatmaps were generated using the deepTools (Ramirez et al., 2014) function computeMatrix with settings ‘reference-point -a 5000 -b 5000’ with spike-in normalised bigWig files and RING1B peaks from Illingworth et al., 2015 merged within 5 kb of each other. Plots were made with the deepTools function plotHeatmap.

### ChIP-qPCR analysis

Dilutions (5-fold) of sonicated input from each mESC sample with MCF-7 spike-in (20% of each IP quantity) were used to create standard curves for the immunoprecipitated H3K27me3 and Ring1B chromatin (2 biological replicates). For each replicate, PCRs were performed in triplicate using LightCycler 480 SYBR Green I detection kit on a BioRad C1000 Touch Thermal Cycler (using primers described in Supplemental Table S10). Thermal cycler programme: 15 min Hotstart; 44 PCR cycles (95°C for 15 s, 55°C for 30 s, 72°C for 30 s).

The qPCR values for H3K27me3 and Ring1B binding to the human *EVX2* gene were used to calibrate the values for polycomb binding at *Hoxd10, En2* and *β-actin* in untreated, 2,5-HD-treated, 1,6-HD-treated and recovered mESCs. The lowest h*EVX2* value for each IP for each replicate was divided by itself and by the other values to generate normalisation factors which were applied to the H3K27me3 and Ring1B binding values for the mouse genes.

### Statistical analysis

DNA-FISH interprobe distance data sets were compared using the two-tailed Mann-Whitney U test, a nonparametric test that compares two unpaired groups. Differences in DNA-FISH data sets comparing categorical distributions were measured using Fisher’s Exact Test. Nuclear size comparisons were measured using Unpaired T-test with Welch’s correction, a parametric test that compares two unpaired groups. All statistical analyses were performed using GraphPad Prism 9.4.1 software (Mann-Whitney, T-test) or online GraphPad 2×2 contingency table www.graphpad.com/quickcalcs/contingency1/ (Fisher’s).

## Data availability

All sequencing data has been submitted to the GEO repository under the accession number GSE224930.

## Acknowledgements

The authors thank the staff of the Institute of Genetics and Cancer imaging facility for their assistance with imaging and the Wellcome Trust Clinical Research Facility (Edinburgh Clinical Research Facility) for sequencing. This work has made use of the resources provided by the Edinburgh Compute and Data Facility (ECDF) (http://www.ecdf.ed.ac.uk/).

## Author contributions

IW conceived and conducted the experiments, analysed the data and wrote the manuscript. SB did the FISH experiments with oligo probe pools, GG and EF helped with bioinformatics analysis, WB conceived the study, secured funding for the work and wrote the manuscript.

