## Supplemental figures and tables for "Dispersal of PRC1 condensates disrupts polycomb chromatin domains and loops"

**Supplementary data**

**Supplemental Figure S1.** Cell morphology and nuclear size of mESCs – biological replicate of the data in Figure 1. (*A*) Representative images of untreated, 2,5-HD-treated, 1,6-HD-treated, and recovered mESCs. Scale bars: 100 μm. Inset scale bars: 20 μm. (*B*) Scatter plot showing the area of each measured nucleus (black dots) for each cell type. Red bars show the mean and the standard deviations. Data sets were tested for differences using the Unpaired T-test with Welch's correction. Data for this figure are in Supplemental Table S1.

**Supplemental Figure S2.** Replicate data for Figure 2 showing reversible chromatin decompaction across polycomb target Hox clusters following 1,6-HD treatment. (*A*) and (*B*) Violin plots show the interprobe distances (nm) for probe pairs shown in Figure 2*A* in untreated, 2,5-HD-treated, 1,6-HD-treated and recovered wild-type cells. (*C*) Violin plots show the interprobe distances (nm) for probe pairs labelling the ends of the HoxB cluster in untreated, 1,6-HD-treated and recovered wild-type cells. * $P \leq 0.05$ and $> 0.01$; ** $P \leq 0.01$; Mann Whitney test. Data for this figure are in Supplemental Table S2.

**Supplemental Figure S3.** Polycomb-dependent associations between *Irx3*, *5*, and *6* and replicate data for Figure 3 showing reversible disruption of the cluster by 1,6-HD. (*A*) Micro-C maps, at 5-kb resolution, of the *Irx3*, *5* and *6* TAD from wild-type (Hsieh et al., 2020), CTCF depleted (CTCF-AID) and cohesin depleted mESCs (RAD21-AID) (Hsieh et al., 2022). Genome co-ordinates (Mb) are from the mm10 assembly of the mouse genome. Genes and RING1B ChIP-seq profiles are shown below. Black arrowheads indicate enriched interactions between *Irx3*, *5*, and *6*. (*B*) Bar plots providing categorical analysis of the spatial location of each of the three polycomb target loci probes shown in Figure 3A relative to each other in untreated, 2,5-HD, 1,6-HD and recovered wild type cells (Biological replicate of Figure 3). Categories are: clustered, at least two of the three loci < 200 nm apart; intermediate, at least two of the three loci between 200 – 399 nm apart; dispersed, all three loci ≥ 400 nm apart. Differences in clustering and dispersal identified using Fisher's Exact test; * P ≤ 0.05 and > 0.01, ** P ≤ 0.01. (*C*) and (*D*) Scatter plots depicting the interprobe distances between each of the two fosmid probe pairs with the separation between the third pair indicated by the color in the color bar in wild-type (*C*) and *Ring1B*$^{-/-}$ cells (*D*). Dashed red box indicates non-dispersed alleles with at least two sets of interprobe distances < 400 nm. Data are summarised in Supplemental Table S3.

**Supplemental Figure S4.** Polycomb-dependent inter-TAD associations between *En2*, *Shh*, and *Mnx1* showing reversible disruption of the associations by 1,6-HD. Replicate data for Figure 4. (*A*) Micro-C maps (5-kb resolution) of the three adjacent TADs on chromosome 5 containing *En2*, *Shh*, and *Mnx1* from wild-type (Hsieh et al., 2020), CTCF-depleted (CTCF-AID) and cohesin depleted mESCs (RAD21-AID) (Hsieh et al., 2022). Genome co-ordinates (Mb) are from the mm10 assembly of the mouse genome. Genes and RING1B ChIP-seq profiles are shown below. Black arrowheads indicate enriched interactions between *En2*, *Shh*, and *Mnx1*. (*B* and *D*) Bar plots providing categorical analysis of the relative spatial location of each of the three polycomb (*B*) and non-polycomb (*D*) target loci probes shown in Figure 4A untreated, 2,5HD, 1,6HD and recovered wild type cells. Biological replicate of the data in Fig. 4. Differences in clustering and dispersal identified using Fisher's Exact test; * P ≤ 0.05 and > 0.01, ** P ≤ 0.01, *** P ≤ 0.001, **** P ≤ 0.0001. (*C, E* and *F)* Scatter plots depicting the interprobe distances between each of the two fosmid probe pairs with the separation between the third pair indicated by the color in the color bar for (*C)* polycomb target loci (replicate for data in Fig. 4D), (*E*) two replicates for non-polycomb target loci in wild-type cells and (*F*) for polycomb target loci in *Ring1B*$^{-/-}$ cells. Dashed red box indicates non-dispersed alleles with at least two sets of interprobe distances < 400 nm. (*G*) Violin plots show the interprobe distances (nm) for probes labelling CTCF binding sites adjacent to *Shh* and the limb enhancer ZRS at the TAD boundaries in untreated, 2,5-HD-treated, 1,6-HD-treated, and recovered wild type cells. Data from two biological replicates. Data for this figure are in Supplemental Table S4.

**Figure S5.** Replicate data for Figure 6 showing reversible reduction of PRC1 binding to polycomb target loci following hexandiol treatment. (*A*) Heat map representation (bottom panel) and summary metaplots (top panel) of RING1B calibrated ChIP-seq signal distribution at RefSeq gene TSS (±5 kb) enriched for marks in wild-type ESCs in untreated, 2,5HD-treated, 1,6HD-treated, and recovered mESCs. (B) UCSC Genome Browser track views of Ring1B cChIP-seq data from untreated, 2,5-HD-treated, 1,6-HD-treated, and recovered mESCs at the HoxD (*top*), *Irx3*, *5*, *6* (*middle*), *En2*, *Shh*, and *Mnx1* (*bottom*) genomic regions. (*C*) As for *B* but for the 50 Mb region of chromosome 2 tiled with oligonucleotide probes.

**Supplemental Table S1. Effects of 2,5 or 1,6 hexanediol on mESC nuclear size.**

| Treatment | Number of nuclei | Nuclear Area ($\mu m^2$) |
|---|---|---|
| **Rep. 1** | | |
| **un** | 50 | 193 |
| **2,5-HD** | 60 | 193 |
| **1,6-HD** | 60 | 194 |
| **rec** | 50 | 193 |
| **Rep. 2** | | |
| **un** | 84 | 167 |
| **2,5-HD** | 86 | 162 |
| **1,6-HD** | 83 | 157 |
| **rec** | 109 | 156 |

Nuclear area of untreated (un) mESCs and mESCs treated with either 2% 2,5 or 1,6 hexandiol (2,5-HD, 1,6-HD) for 5 minutes and for cells > 1-hour post-1,6-HD treatment (rec). Data are from two biological replicates (Rep.1 and Rep.2). Areas were measured by flattening (extended focus) the deconvolved 3D DAPI-stained images and calculating nuclear area using the freehand ROI tool on Volocity (see methods). Nuclear areas indicated ($\mu m^2$) are mean values, statistical analysis by unpaired T Tests (two-tailed).

**Supplemental Table S2. Effects of 2,5 or 1,6 hexanediol on chromatin compaction at the HoxD and HoxB loci**

| Treatment | GCR-Lnp | Evx2-Hoxd3 | Evx2-Hoxd3 (*Ring1B*$^{-/-}$) | Hoxb13-Hoxb1 |
|---|---|---|---|---|
| | Interprobe distance (nm) and number of alleles [ ] | | | |
| **Rep. 1** | | | | |
| **un** | 329 [122] | 272 [92] | 420 [82] | 494 [149] |
| **2,5-HD** | 334 [112] | 262 [100] | | |
| **1,6-HD** | 288 [123] | 326 ($p$ = 0.0099) [79] | 379 [81] | 515 ($p$ = 0.038) [119] |
| **rec** | 287 [126] | 268 [88] | 379 [60] | 496 [169] |
| **Rep. 2** | | | | |
| **un** | 327 [100] | 307 [169] | | |
| **2,5-HD** | 271 [100] | 286 [128] | | |
| **1,6-HD** | 288 [90] | 333 ($p$ = 0.019) [130] | | |
| **rec** | 297 [86] | 323 [132] | | |

Statistical analysis of data for Fig. 2C, E, G & Supplemental Fig. S2A, B, C. Interprobe distances measured across the HoxB (Hoxb13-Hoxb1) and HoxD (Evx2-Hoxd3) loci and at a control locus adjacent to HoxD (GCR-Lnp) in two biological replicates of untreated (un) mESCs and in cells treated with 2% 2,5 or 1,6 hexanediol and for cells > 1-hour post-1,6-HD treatment (rec). Data from mESCs mutant for PRC1 (*Ring1B*$^{-/-}$) are also shown. Interprobes distances shown are median values in nm. Square brackets indicate the number of alleles measured. $p$-values <0.05 from Mann-Whitney U Tests are indicated.

**Supplemental Table S3. Effects of 2,5 or 1,6 hexanediol on clustering of *Irx3*, *Irx5*, and *Irx6***

| Treatment | Wild type | Ring1B$^{-/-}$ |
|---|---|---|
| | Clustering (≤ 200 nm) frequency (%) of minimum of 2 of 3 loci and number of alleles [ ] | |
| **Rep. 1** | | |
| **un** | 23 [198] | 10 [108] |
| **2,5-HD** | 25 [133] | |
| **1,6-HD** | 15 ($p$ = 0.019) [178] | 13 [136] |
| **rec** | 30 [165] | 7 [126] |
| **Rep. 2** | | |
| **un** | 26 [103] | |
| **2,5-HD** | 21 [135] | |
| **1,6-HD** | 8 ($p$ = 0.002) [133] | |
| **rec** | 21 [122] | |
| | Dispersed (≥ 400 nm) frequency (%) of all 3 loci | |
| **Rep. 1** | | |
| **un** | 19 | 30 |
| **2,5-HD** | 16 | |
| **1,6-HD** | 38 ($p$ = 0.0002) | 35 |
| **rec** | 14 | 31 |
| **Rep. 2** | | |
| **un** | 19 | |
| **2,5-HD** | 25 | |
| **1,6-HD** | 37 ($p$ = 0.02) | |
| **rec** | 26 | |

Statistical analysis of data for Figs. 3C, F; Supplementary Fig. S3B. The proportion of alleles with clustering of *Irx3*, *Irx5*, and *Irx6* (≤ 200 nm between at least 2 of the three loci) or with dispersed alleles (≥ 400 nm between all 3 loci) in untreated (un) wild-type mESCs and in cells treated with 2% 2,5 or 1,6 hexanediol and for cells > 1-hour post-1,6-HD treatment (rec). Data from mESCs mutant for PRC1/Ring1B (R1B$^{-/-}$) are also shown. *p*-values from Fisher's Exact Tests. Data are from two independent biological replicates.

**Table S4. The proportion of *En2*, *Shh*, and *Mnx1* clustering and dispersed alleles in 1,6-HD-treated wild type and R1B$^{-/-}$ mESCs compared to un, 2,5-HD and rec**

| Treatment | Wild type mESCs | R1B$^{-/-}$ mESCs |
|---|---|---|
| | **Clustering (≤ 200 nm) frequency (%) of minimum of 2 polycomb targets and number of alleles [ ]** | |
| **Rep. 1** | | |
| **un** | 34 [100] | 7 [103] |
| **2,5-HD** | 29 [111] | |
| **1,6-HD** | 13 ($p$ = 0.0007) [100] | 13 [90] |
| **rec** | 41 [100] | 11 [90] |
| **Rep. 2** | | |
| **un** | 27 [100] | |
| **2,5-HD** | 24 [100] | |
| **1,6-HD** | 6 ($p$ < 0.0001) [100] | |
| **rec** | 20 [100] | |
| | **Dispersed (≥ 400 nm) frequency (%) of all 3 polycomb targets** | |
| **Rep. 1** | | |
| **un** | 7 | 46 |
| **2,5-HD** | 12 | |
| **1,6-HD** | 32 ($p$ < 0.0001) | 34 |
| **rec** | 10 | 41 |
| **Rep. 2** | | |
| **un** | 19 | |
| **2,5HD** | 17 | |
| **1,6HD** | 34 ($p$ = 0.02) | |
| **rec** | 24 | |

Statistical analysis of data for Figs. 4C, F & S4A. *p*-values from Fisher's Exact Tests.

**Table S5. The proportion of *Cnpy1*, SFPE1, and *Ube3c* clustering and dispersed alleles in 1,6-HD-treated wild type mESCs compared to un, 2,5-HD and rec**

| Treatment | Wild type mESCs |
|---|---|
| | Clustering (≤ 200 nm) frequency (%) of minimum of 2 non-polycomb targets and number of alleles [ ] |
| **Rep. 1** | |
| **un** | 15 [150] |
| **2,5-HD** | 7 [60] |
| **1,6-HD** | 10 [100] |
| **rec** | 6 [84] |
| **Rep. 2** | |
| **un** | 4 [91] |
| **2,5-HD** | 10 [75] |
| **1,6-HD** | 7 [61] |
| **rec** | 10 [61] |
| | Dispersed (≥ 400 nm) frequency (%) of all 3 non-polycomb targets |
| **Rep. 1** | |
| **un** | 32 |
| **2,5-HD** | 42 |
| **1,6-HD** | 36 |
| **rec** | 27 |
| **Rep. 2** | |
| **un** | 43 |
| **2,5-HD** | 39 |
| **1,6-HD** | 31 |
| **rec** | 28 |

Statistical analysis of data for Figs. 4E & S4B. $p$-values from Fisher's Exact Tests.

**Table S6. Comparison of the number of discrete polycomb and non-polycomb target foci in untreated mESCs with 2,5HD-treated, 1,6-HD-treated and recovered mESCs across chromosome 2**

| Treatment | Polycomb + | Polycomb - |
|---|---|---|
| | Number of foci and number of alleles [ ] | |
| **un** | 15 [119] | 16 [124] |
| **2,5-HD** | 16 ($p$ = 0.02) [122] | 18 ($p$ = 0.007) [125] |
| **1,6-HD** | 16 ($p$ = 0.001) [123] | 16 [127] |
| **rec** | 13 ($p$ = 0.047) [158] | 13 ($p$ < 0.0001) [155] |

Statistical analysis of data for Fig. 5C. Foci numbers are median values, $p$-values from Mann-Whitney U Tests.

**Table S7. Chr2 MyTags H3K27me3+ Regions**

| Chromosome | Start | End | Ring1B peak |
|---|---|---|---|
| chr2 | 42498000 | 42518000 | 1 |
| chr2 | 44380000 | 44400000 | 1 |
| chr2 | 49633000 | 49653000 | 1 |
| chr2 | 52521000 | 52541000 | 1 |
| chr2 | 54278000 | 54298000 | 0 |
| chr2 | 55278000 | 55298000 | 1 |
| chr2 | 56951000 | 56971000 | 1 |
| chr2 | 58200000 | 58220000 | 1 |
| chr2 | 60113000 | 60133000 | 1 |
| chr2 | 61640000 | 61660000 | 1 |
| chr2 | 65068000 | 65088000 | 0 |
| chr2 | 65952000 | 65972000 | 1 |
| chr2 | 68297000 | 68317000 | 1 |
| chr2 | 69410000 | 69430000 | 1 |
| chr2 | 70393000 | 70413000 | 1 |
| chr2 | 71365000 | 71385000 | 1 |
| chr2 | 73100000 | 73120000 | 1 |
| chr2 | 74560000 | 74580000 | 1 |
| chr2 | 76233000 | 76253000 | 1 |
| chr2 | 77644000 | 77664000 | 1 |
| chr2 | 79086000 | 79106000 | 0 |
| chr2 | 80276000 | 80296000 | 1 |
| chr2 | 81883000 | 81903000 | 1 |
| chr2 | 84705000 | 84725000 | 1 |
| chr2 | 90717000 | 90737000 | 1 |
| chr2 | 91754000 | 91774000 | 1 |
| chr2 | 92745000 | 92765000 | 0 |
| chr2 | 93785000 | 93805000 | 1 |

Mouse genome assembly number: NCBI m37. Probes previously used in (Boyle et al., 2020).

**Table S8. Chr2 MyTags H3K27me3- Regions**

| Chromosome | Start | End |
|---|---|---|
| chr2 | 43120000 | 43140000 |
| chr2 | 46675000 | 46695000 |
| chr2 | 51945000 | 51965000 |
| chr2 | 53038000 | 53058000 |
| chr2 | 56165000 | 56185000 |
| chr2 | 57571000 | 57591000 |
| chr2 | 59077000 | 59097000 |
| chr2 | 60673000 | 60693000 |
| chr2 | 63837000 | 63857000 |
| chr2 | 65285000 | 65305000 |
| chr2 | 67600000 | 67620000 |
| chr2 | 68800000 | 68820000 |
| chr2 | 69717000 | 69737000 |
| chr2 | 70862000 | 70882000 |
| chr2 | 72210000 | 72230000 |
| chr2 | 73524000 | 73544000 |
| chr2 | 75490000 | 75510000 |
| chr2 | 77056000 | 77076000 |
| chr2 | 78762000 | 78782000 |
| chr2 | 79510000 | 79530000 |
| chr2 | 81255000 | 81275000 |
| chr2 | 82760000 | 82780000 |
| chr2 | 84272000 | 84292000 |
| chr2 | 88200000 | 88220000 |
| chr2 | 91220000 | 91240000 |
| chr2 | 92090000 | 92110000 |
| chr2 | 93580000 | 93600000 |
| chr2 | 94233000 | 94253000 |

Mouse genome assembly number: NCBI m37. Probes previously used in (Boyle et al., 2020).

### Table S9. Fosmid Probes

| | Region | Whitehead (Sanger) Name | Ensemble name | Coordinates Start | End | Size (bp) |
|---|---|---|---|---|---|---|
| **Hoxd** | *GCR** | WI1-2157A11 | G135P63331H7 | 74242615 | 74282044 | 39429 |
| | *Lnp** | WI1-482L15 | G135P61870C5 | 74329582 | 74372986 | 43404 |
| | *Evx2-Hoxd13** | WI1-469P2 | G135P67444A12 | 74474157 | 74513003 | 38846 |
| | *Hoxd4-Hoxd1** | WI1-121N10 | G135P67844B8 | 74566983 | 74605438 | 38455 |
| **Irx** | *Irx3*** | WI1-1206E19 | | 94307305 | 94341470 | 34166 |
| | *Irx5*** | WI1-1060D04 | | 94865335 | 94904882 | 39548 |
| | *Irx6*** | WI1-0901O22 | | 95190971 | 95221152 | 30182 |
| **Shh** | *En2**** | WI1-2728F4 | | 28477913 | 28517563 | 39650 |
| | *Cnpy1**** | WI1-2816N11 | | 28567538 | 28605641 | 38104 |
| | *Shh**** | WI1-574O18 | G135P64333A4 | 28754458 | 28795879 | 41421 |
| | *SFPE1**** | WI1-0552F05 | | 28798551 | 28841224 | 42673 |
| | *ZRS***** | WI1-1047E14 | G135P600929F6 | 29611727 | 29653695 | 41968 |
| | *Mnx1**** | WI1-1204B6 | | 29791124 | 29827491 | 36368 |
| | *Ube3c**** | WI1-0492O10 | | 29873876 | 29916051 | 42186 |

Names are Ensembl (r 45) (http://jun2007.archive.ensembl.org/Mus_musculus/index.html). Mouse genome assembly number: NCBI m37. Asterisks indicate fosmids previously used in: * (Williamson et al., 2012) ** (Sobreira et al., 2021) *** (Boyle et al., 2020) **** (Williamson et al., 2016, 2019)

### Table S10. qPCR primers for ChIP analysis

| Promoter/exon | Oligo name | Sequence |
|---|---|---|
| *Hoxd10* | Hoxd10prof | TAGTAGATGTCGCTGTTGTCCG |
| | Hoxd10pror | ACATGACAACCAAGCCAATGAGA |
| *En2* | En2intronf | CAACTCTGGGTGCTCTCCTG |
| | En2intronr | GCTTGCAGGATGGAACGAAC |
| *Actin* | Actinf | CCTCGATGCTGACCCTCATCC |
| | Actinr | GACACTGCCCCATTCAATGTCTC |
| h*EVX2* | EVX2prof | GAGACAAGTGAGGAGGAA |
| | EVX2pror | AACTGAGAGCGTAATGATG |

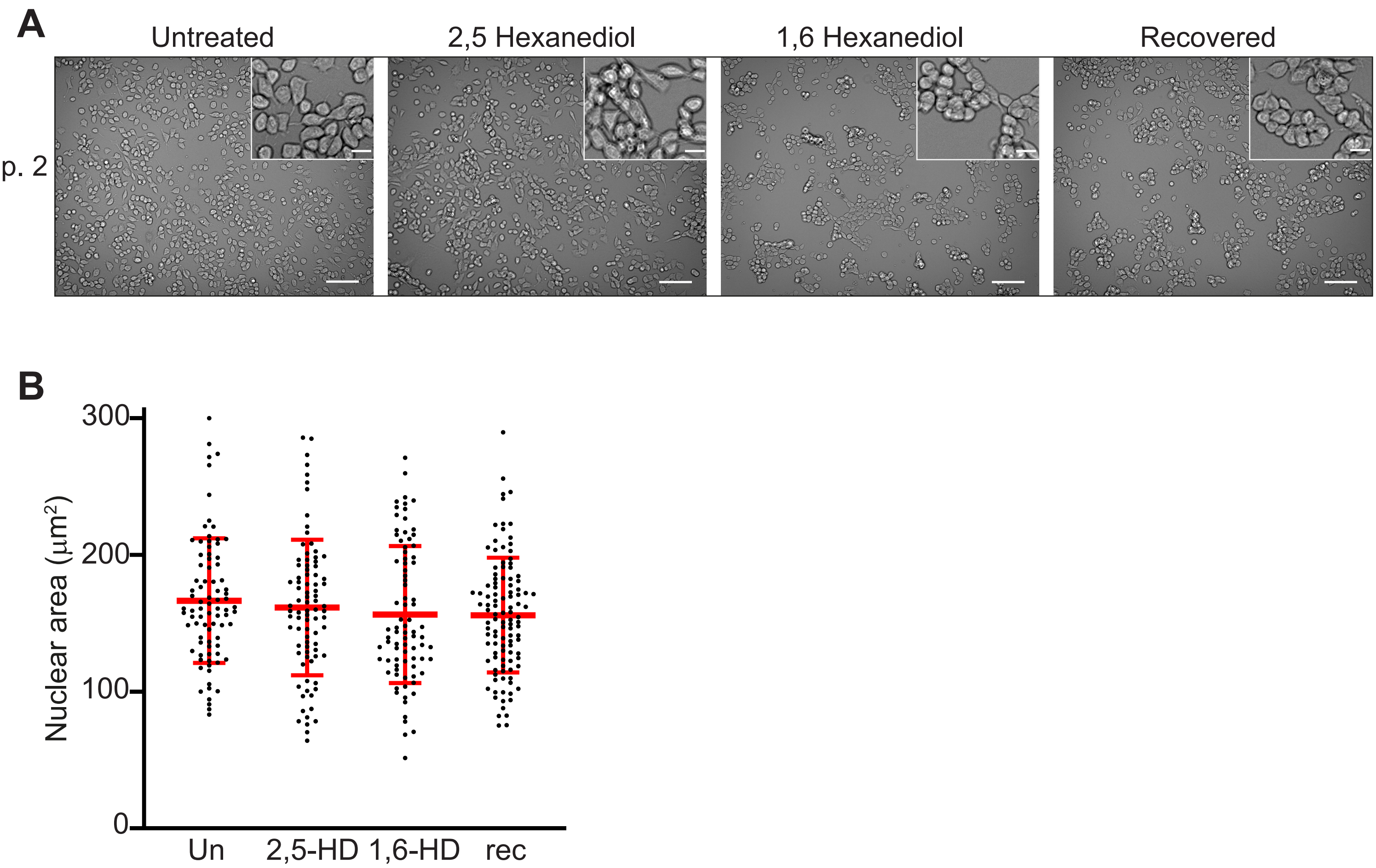


**Supplemental Figure 1.** Cell morphology and nuclear size of mESCs – biological replicate of the data in Figure 1. (*A*) Representative images of untreated, 2,5-HD-treated, 1,6-HD-treated, and recovered mESCs. Scale bars: 100 µm. Inset scale bars: 20 µm. (*B*) Scatter plot showing the area of each measured nucleus (black dots) for each cell type. Red bars show the mean and the standard deviations. Data sets were tested for differences using the Unpaired T-test with Welch's correction. Data for this figure are in Supplemental Table S1.

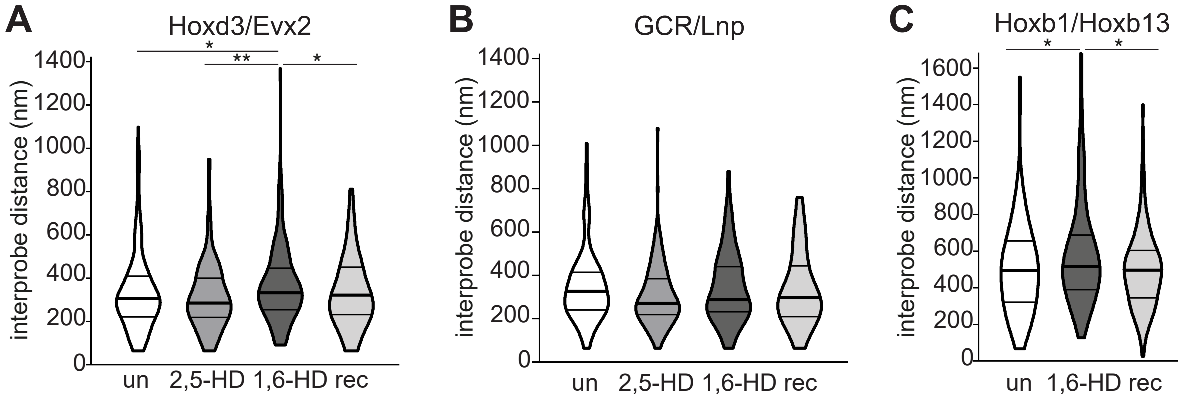


**Supplemental Figure 2.** Replicate data for Figure 2 showing reversible chromatin decompaction across polycomb target Hox clusters following 1,6-HD treatment. (*A*) and (*B*) Violin plots show the interprobe distances (nm) for probe pairs shown in Figure 2*A* in untreated, 2,5-HD-treated, 1,6-HD-treated and recovered wild-type cells. (*C*) Violin plots show the interprobe distances (nm) for probe pairs labelling the ends of the HoxB cluster in untreated, 1,6-HD-treated and recovered wild-type cells. * $P \leq 0.05$ and $> 0.01$; ** $P \leq 0.01$; Mann Whitney test. Data for this figure are in Supplemental Table S2.

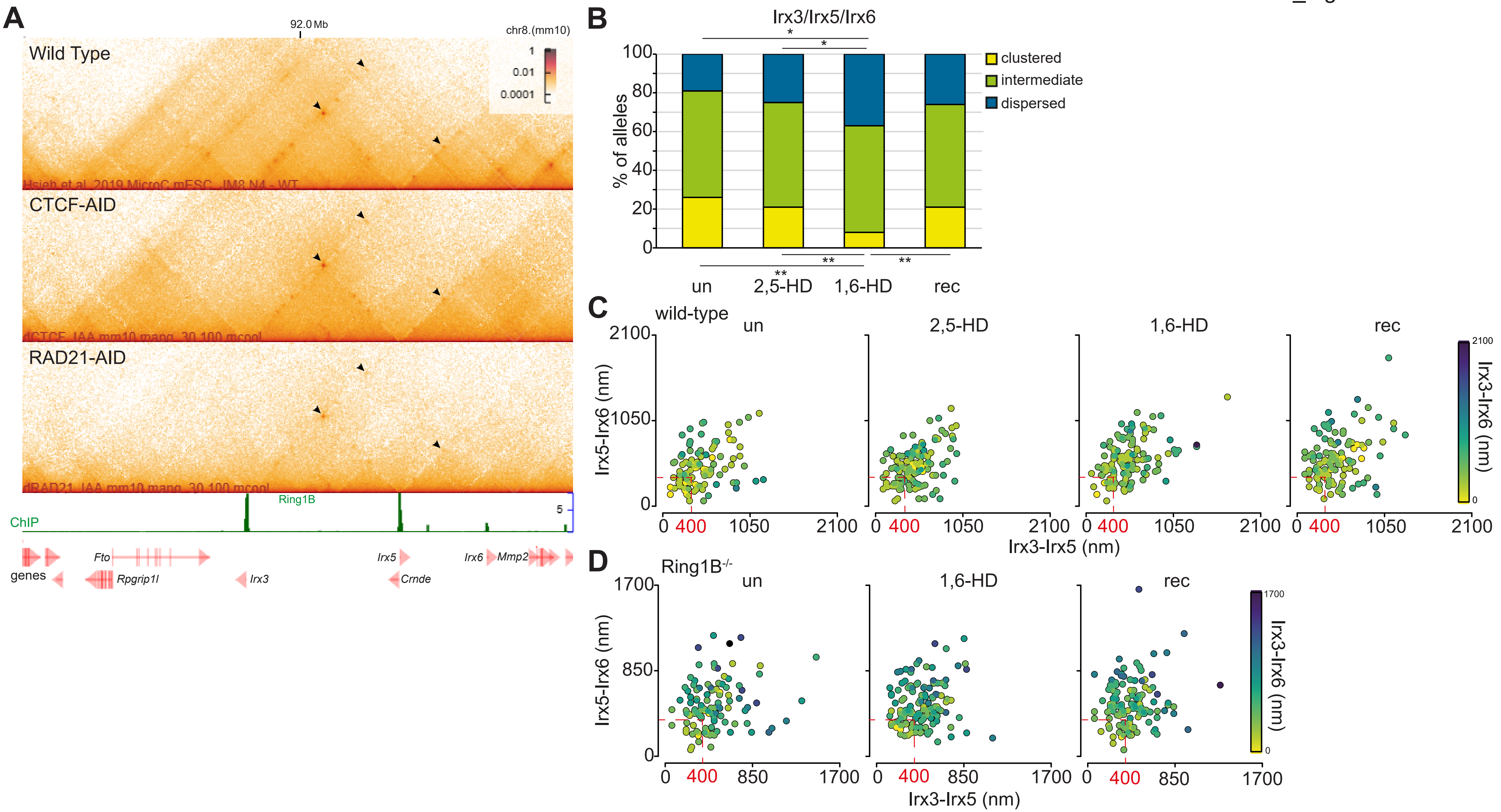


**Figure S3.** Polycomb-dependent associations between Irx3, 5, and 6 and replicate data for Figure 3 showing reversible disruption of the cluster by 1,6-HD. (*A*) Micro-C maps, at 5-kb resolution, of the *Irx3*, *5* and *6* TAD from wild-type (Hsieh et al., 2020), CTCF depleted (CTCF-AID) and cohesin depleted mESCs (RAD21-AID) (Hsieh et al., 2022). Genome co-ordinates (Mb) are from the mm10 assembly of the mouse genome. Genes and RING1B ChIP-seq profiles are shown below. Black arrowheads indicate enriched interactions between *Irx3*, *5*, and *6*. (*B*) Bar plots providing categorical analysis of the spatial location of each of the three polycomb target loci probes shown in Figure 3*A* relative to each other in untreated, 2,5-HD, 1,6-HD and recovered wild type cells (Biological replicate of Figure 3). Categories are: clustered, at least two of the three loci < 200 nm apart; intermediate, at least two of the three loci between 200 – 399 nm apart; dispersed, all three loci ≥ 400 nm apart. Differences in clustering and dispersal identified using Fisher's Exact test; * P ≤ 0.05 and > 0.01, ** P ≤ 0.01. (*C*) and (*D*) Scatter plots depicting the interprobe distances between each of the two fosmid probe pairs with the separation between the third pair indicated by the color in the color bar in wild-type (*C*) and *Ring1B-/-* cells (*D*). Dashed red box indicates non-dispersed alleles with at least two sets of interprobe distances < 400 nm. Data are summarised in Supplemental Table S3.

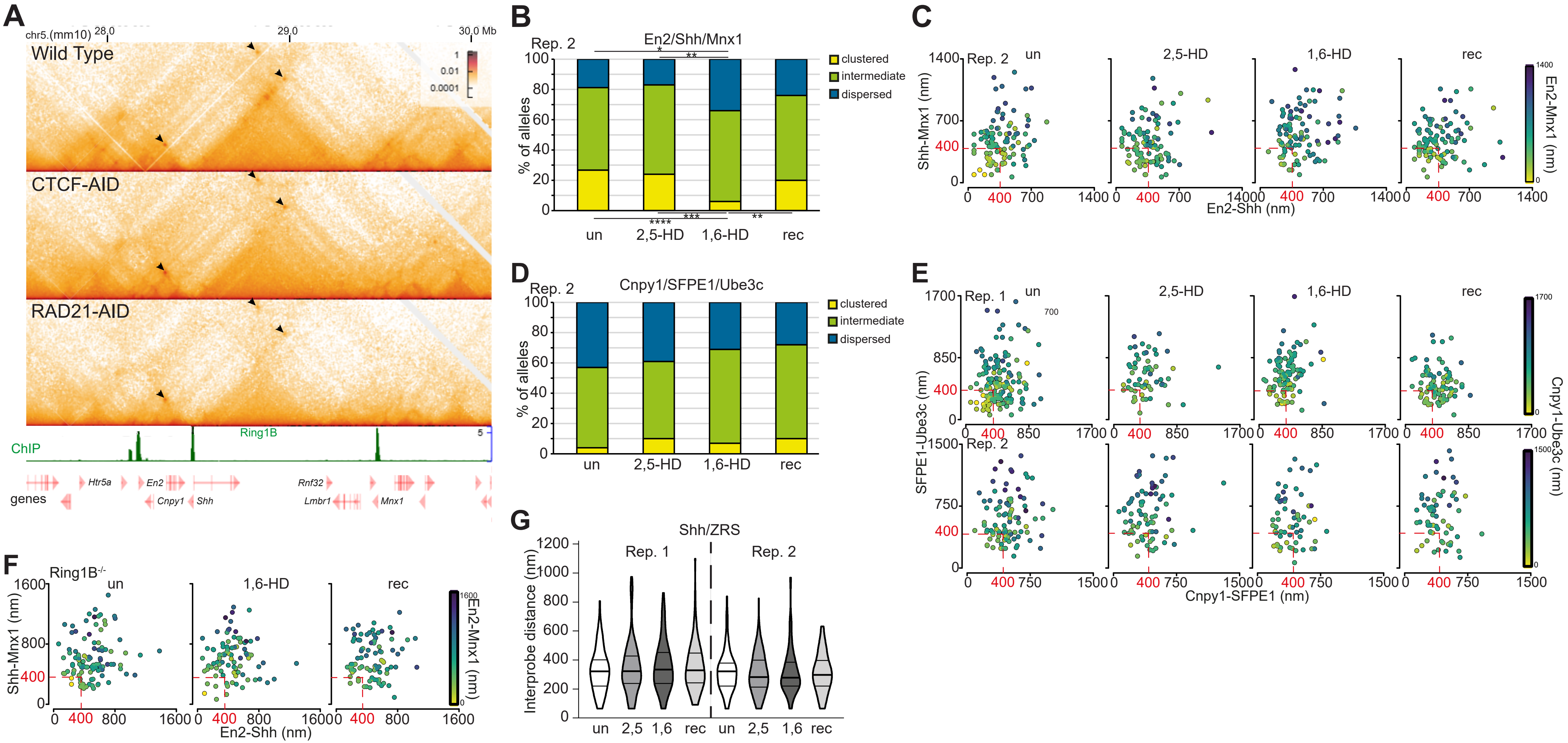


**Figure S4.** Polycomb-dependent inter-TAD associations between *En2*, *Shh*, and *Mnx1* showing reversible disruption of the associations by 1,6-HD. Replicate data for Figure 4. (*A*) Micro-C maps (5-kb resolution) of the three adjacent TADs on chromosome 5 containing *En2*, *Shh*, and *Mnx1* from wild-type (Hsieh et al., 2020), CTCF-depleted (CTCF-AID) and cohesin depleted mESCs (RAD21-AID) (Hsieh et al., 2022). Genome co-ordinates (Mb) are from the mm10 assembly of the mouse genome. Genes and RING1B ChIP-seq profiles are shown below. Black arrowheads indicate enriched interactions between *En2*, *Shh*, and *Mnx1*. (*B* and *D*) Bar plots providing categorical analysis of the relative spatial location of each of the three polycomb (*B*) and non-polycomb (*D*) target loci probes shown in Figure 4*A* untreated, 2,5HD, 1,6HD and recovered wild type cells. Biological replicate of the data in Fig. 4. Differences in clustering and dispersal identified using Fisher's Exact test; * $P \leq 0.05$ and $> 0.01$, ** $P \leq 0.01$, *** $P \leq 0.001$, **** $P \leq 0.0001$. (*C*, *E* and *F*) Scatter plots depicting the interprobe distances between each of the two fosmid probe pairs with the separation between the third pair indicated by the color in the color bar for (*C*) polycomb target loci (replicate for data in Fig. 4*D*), (*E*) two replicates for non-polycomb target loci in wild-type cells and (*F*) for polycomb target loci in *Ring1B-/-* cells. Dashed red box indicates non-dispersed alleles with at least two sets of interprobe distances < 400 nm. (*G*) Violin plots show the interprobe distances (nm) for probes labelling CTCF binding sites adjacent to *Shh* and the limb enhancer ZRS at the TAD boundaries in untreated, 2,5-HD-treated, 1,6-HD-treated, and recovered wild type cells. Data from two biological replicates. Data for this figure are in Supplemental Table S4.

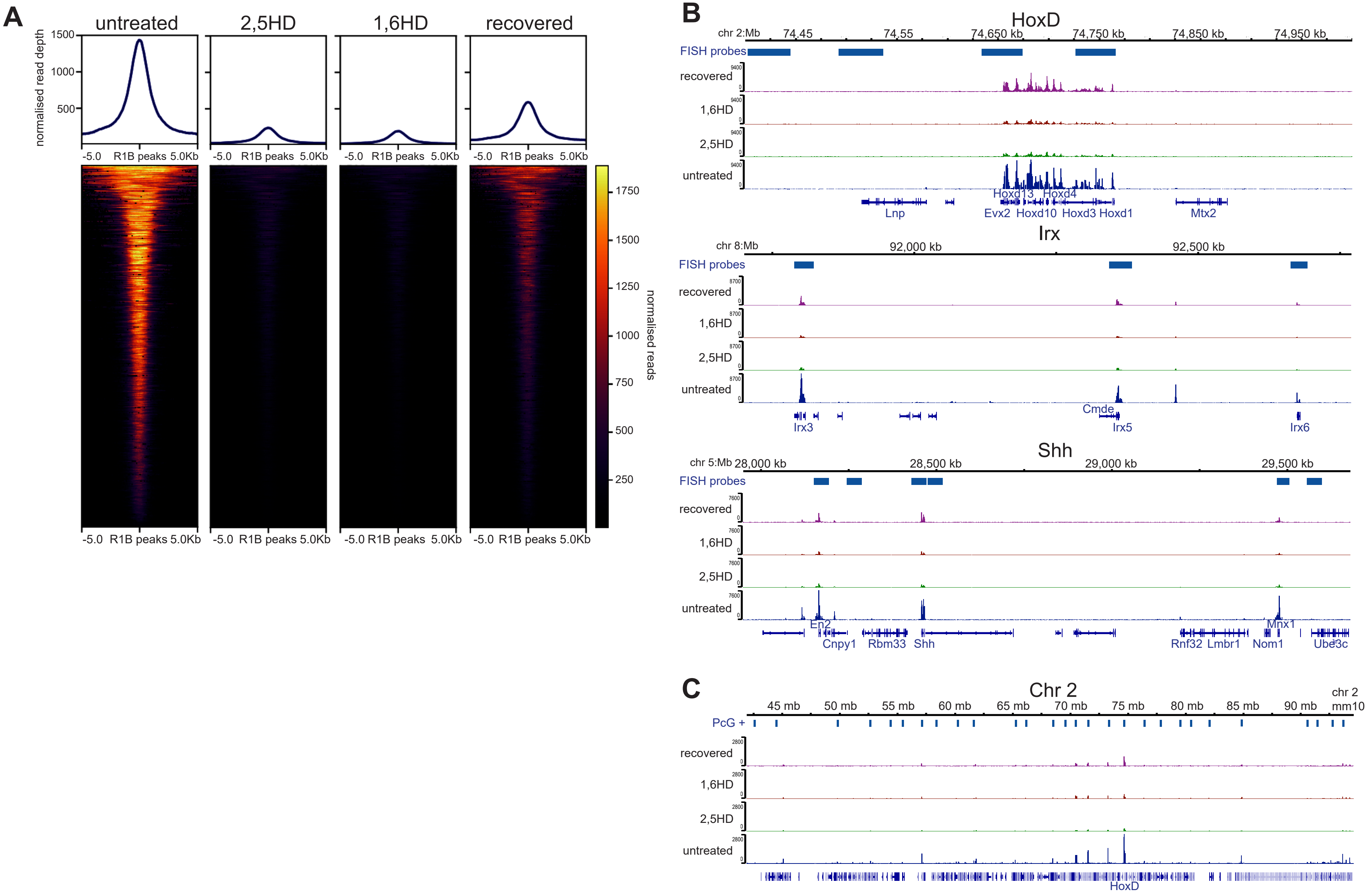


**Figure S5.** Replicate data for Figure 6 showing reversible reduction of PRC1 binding to polycomb target loci following hexandiol treatment. (*A*) Heat map representation (bottom panel) and summary metaplots (top panel) of RING1B calibrated ChIP-seq signal distribution at RefSeq gene TSS (±5 kb) enriched for marks in wild-type ESCs in untreated, 2,5HD-treated, 1,6HD-treated, and recovered mESCs. (*B*) UCSC Genome Browser track views of RING1B cChIP-seq data from untreated, 2,5-HD-treated, 1,6-HD-treated, and recovered mESCs at the HoxD (top), *Irx3*, *5*, *6* (middle), *En2*, *Shh*, and *Mnx1* (bottom) genomic regions. (*C*) As for B but for the 50 Mb region of chromosome 2 tiled with oligonucleotide probes.
